# Small molecule promoters of endogenous lipid droplet accumulation drive lysophagy

**DOI:** 10.1101/2025.01.10.632143

**Authors:** Sai Kumari Vechalapu, Santhosh Duraisamy, Sathyapriya Senthil, Rakesh Kumar, Deepa Ajnar, Apparao Draksharapu, Suresh Kumar, Dharmaraja Allimuthu

## Abstract

Lipid droplets (LDs) play a central role in regulating metabolism in stress-induced conditions, including one triggered by nutrient deprivation. Unravelling the protein networks involved in the biogenesis of LDs and their causative and functional roles in health and disease continue to evolve. To this cause, genetic manipulation of the lipid metabolic network or supplementation of high fat diet/ oleic acid (OA) are the traditional routes for voluntarily triggering LDs formation in cells and animals. We developed a screening platform for the identification of new LDs inducers, and our primary screening of various fatty acids identified linoleic acid (LOA, DUFA) as a better tool than OA (MUFA) in promoting LDs formation. The screening and validation discovered new small molecule-based tools for promoting a rapid organization of endogenous lipids into droplets in multiple cell types. Notably, our mass spectral lipidomics analysis presented the overproduction of phosphatidylcholines and small triglycerides, a hallmark of LDs. Mechanistic investigations of our lead molecules highlighted lipid peroxidation and ATP depletion through mitochondrial impairment in cells, which could serve as chemical cues for driving the fusion of cellular lipids into LDs. Finally, we uncovered the abrupt levels of LDs formation induced by our molecules promoted lysophagy in cancer cells to prevent their proliferation. Collectively, our work introduces new small molecules as powerful tools for reliably promoting LDs accumulation for studying their roles in biology, and we demonstrate the over accumulation of LDs prevent cancer cell proliferation, movement, and colonization.

## Introduction

Lipid droplets (LDs) composed of neutral triglycerides (TG) are wrapped with a phospholipid monolayer are cytoplasmic organelles that serve as crucial reservoirs for energy storage and release fatty acids through lipolysis during metabolic demand.^1–8^ The orchestration of lipid accumulation into droplet is driven by several proteins at the endoplasmic reticulum.^1–5,9^ Eukaryotic cells universally package lipids into lipid droplets in response to chemical metabolic cues received from nutrient deprived states of cells and tissues.^1,2,10^ Beyond energy homeostasis, LDs regulate lipid metabolism, sequester excess or toxic lipids, and play vital roles in cellular signaling, protein degradation, and stress responses.^10,11^ Their functional versatility impacts diverse processes, including membrane biogenesis, oxidative stress protection, and pathogen defense, underscoring their importance in maintaining cellular and organismal health.^1,2,10,11^ Particularly, fatty acids are delivered from LDs to mitochondria for β-oxidation to generate energy during starvation or nutrient stress conditions,^12–15^ except in brown adipocytes.^16^ Understanding the biogenesis of LDs and its biological significance in various cellular events is of high interest.^1,2,10^

Given the importance of LDs roles in biology, a controlled induction of endogenous lipid droplet accumulation in cells using small molecules could facilitate the understanding of precise roles of endogenous LDs and to investigate their mechanism of action. Traditionally, LDs are induced by exposing cells or animals to high fat diet or oleic acid (OA) for longer durations at high micromolar (100s) concentrations, or through genetic manipulations of genes involved in lipid metabolism, such as ATGL.^17–22^ The usage of FA in high doses for cell-base and *in vivo* experiments are associated with their poor solubility and limited cell permeability, otherwise high fat diet is the only alternative. Interestingly, non-fatty acid class of small molecules for a regulated induction of LDs in cells are not common. Endogenous lipids tend to accumulate into LDs as an alternate source for energy production during starvation to counter the energy demand.^1,11^ In parallel, lipids organize themselves into a droplet as a protective mechanism from ROS during enhanced lipid oxidative situations.^11,23,24^ Recent findings highlights the involvement of lipid droplets in preventing lipid peroxidation, thereby control ferroptosis.^25,26^ We hypothesized that small molecules capable of promoting lipid peroxidation has the capacity to trigger LDs formation in cells. Here, we developed a phenotypic screening using high-content imaging platform for studying natural fatty acids to single out the most efficient FA and new small molecules as LD inducers in hepatocellular carcinoma (Huh-7) cells. We report LOA and a set of small molecules that rapidly induce lipid droplets at significantly lower concentrations than OA and other fatty acids in multiple cell lines. The hallmark of LDs was characterized by mass spectral lipidomics and combination experiments. We demonstrated that the lead small molecules target mitochondrial function and trigger lipid peroxidation, thereby promoting the LDs formation. Finally, the accumulation of unusual levels of LDs was observed to drive lysosomal damage, a mechanism called lysophagy in cancer cells to prevent their proliferation at sub-micromolar concentrations and inhibits their migration and colonization.

## Results

### High-content imaging identifies LOA was identified as an optimal fatty acid (FA) for LDs induction among fatty acids

For the identification of chemical-based lipid droplet inducers in cells, we have developed an imaging-based phenotypic screening platform capturing lipid droplets stained with Lipi green (LipiG) fluorophore (Figure 1A).^27^ Aggravated lipid metabolism is a hallmark of hepatocellular carcinoma cells; therefore, Huh-7 cells were used as a model system in our study.^19^ The assay system was established by exposing Huh-7 cells to oleic acid (OA) as positive control and DMSO (0.01%, v/v) as a vehicle control. The hit molecules identified in our screen induced LDs formation greater than three standard deviations (z-score) on DMSO control, which was comparable to that of OA at 200 µM dose.

**Figure 1.**
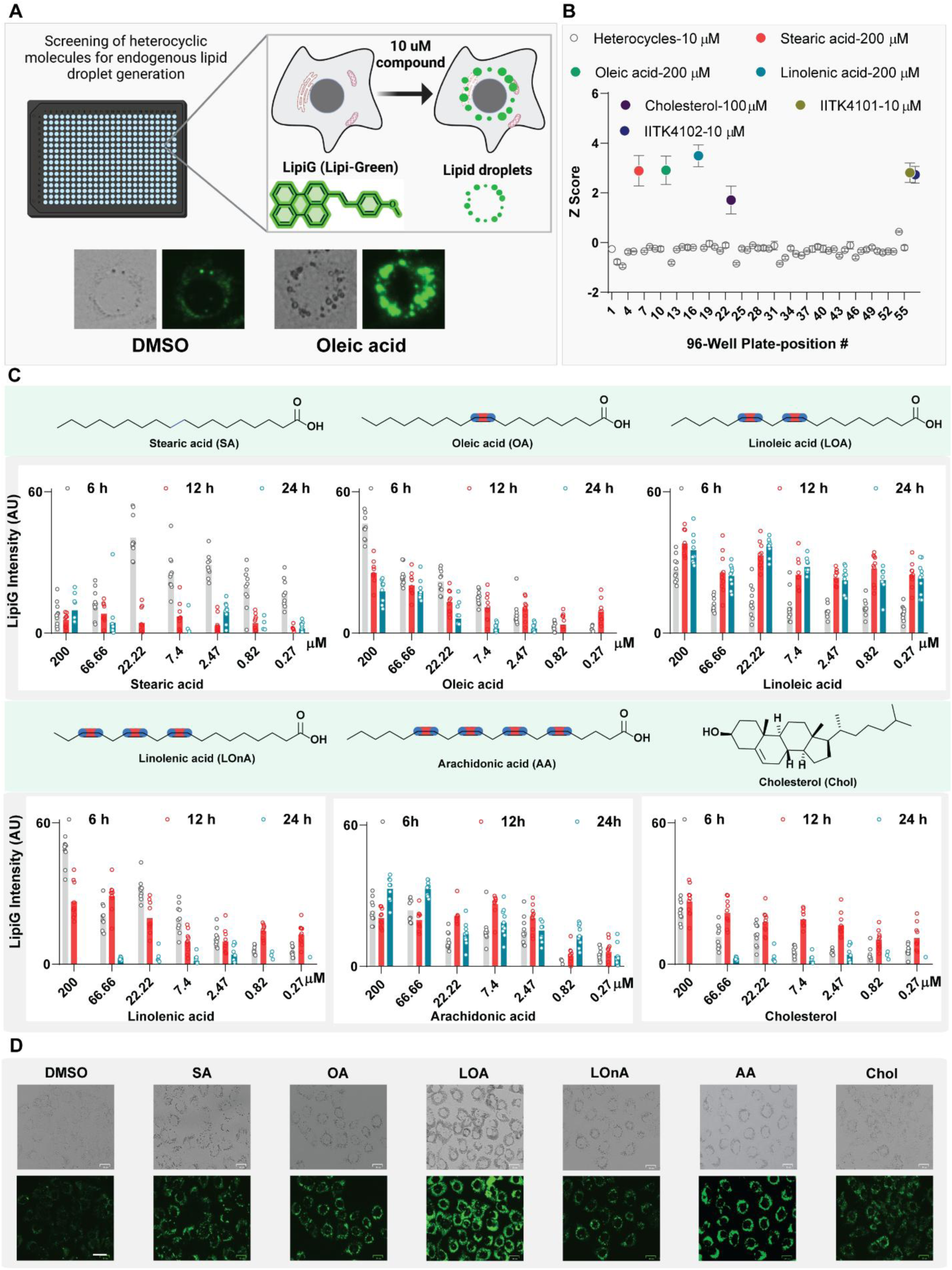
Phenotypic screening and systematic investigation of fatty acids identifies LOA potent inducer of LDs formation in Huh-7 cells. (A) A schematic representation of lipid droplet imaging in cells in our high-content imaging platform and the representative images of Huh-7 cells exposed to DMSO and OA (at 200 µM) and stained with Lipi green for LDs. (B). Screening data of a small library of heterocyclic small molecules at 10 µM (57) and fatty acids plotted z-score for LDs quantification against DMSO control. (C) LDs induced by indicated fatty acids quantified in does and time-dependent fashion in Huh-7 cells. (D)Representative images of images of Huh-7 cells treated with 50 µM of Fatty acids and vehicle control (DMSO), Scale bar: 25 µm.

Here, we set out to investigate the utility of our phenotypic screening platform for the investigation of optimal number of unsaturation sites essential in the FA structure to promote LDs formation in cells. For this study, we assembled saturated FA (stearic acid: C18:0, SA), monounsaturated (MUFA, oleic acid: C18:1 Δ^9^, OA) and diunsaturated fatty acid (DUFA, linoleic acid: C18:2 Δ^9,12^), and polyunsaturated fatty acids (LOA; linolenic acid: C18:3 Δ^9, 12, 15^, LOnA; and arachidonic acid: C20:4 Δ^5, 8, 11, 14^, AA), along with cholesterol, a non-fatty acid lipid (Figure 1A, B). Interestingly, among fatty acids included in the primary screening, LOA and SA were found to enhance LDs formation in huh-7 cells and the same was not true for cholesterol at comparable doses (Figure 1B). Subsequent validation with dose and time-dependent analysis of all these FAs, starting from 200 µM, in a 7-point dose ending at 270 nM in Huh-7 cells revealed a maximal effect for LOA. While the lipid staining was observed to be higher for SA within 6 h at various concentrations, it was noted to be highly toxic to cells at treatment doses above 7.4 µM (Figure 1C, D, Extended data figure 1A). MUFA and DUFA were found to be nontoxic at the highest concentration tested (200 µM), however, a trend of increasing the number of double bonds beyond 2 as in PUFAs (LOnA (3) and AA (4)) were noted to be toxic at higher concentrations (Figure 1C, D, Extended data figure 1A). In the case of cholesterol, neither toxicity nor LDs accumulation was recorded in the tested conditions (Figure 1C, D, Extended data figure 1A). Overall, using our screening platform, the effort towards the identification of optimal DBs needed for an effective induction of LDs formation revealed LOA (DUFA) as a better LDs inducer than MUFA (OA) under comparable conditions.

### Our screening platform identified new small molecule inducers of lipid droplets

Our primary screening of small molecule collections (10 µM) has identified IITK4101 and its iron complex (IITK4102) as the top hits (Figure 1 A, B). We revalidated the screening results hit molecules IITK4101 and IITK4102 in a dose-dependent fashion for reproducibility (Figure 2A, B). The investigation of spatiotemporal effect of our molecules in LDs formation indicated the lead compounds are capable of inducing LDs at as low as 50 nM concentration in 2 h, and the maximum effect was recorded for 410 nM dose after 2 h of treatment for both molecules (Figure 2A-D). The cells are expected to respond to the accumulation of LDs through the overexpression of proteins involved in disrupting the LDs.^28,29^ The hydrolysis of triglycerides by ATGL (adipose triglyceride lipase) is a natural cellular response in conditions were LDs are overproduced.^28^ We performed immunoblotting analysis for ATGL in Huh-7 cells proteome exposed to IITK4101 and IITK4102 in a dose and time dependent fashion. We observed a clear upregulation of ATGL proteins after 6 h of the compound treated conditions and the effect was found to be moderate in time-dependent analysis (Figure 2E). LDs possess triglycerides and cholesterol esters at the core which are enveloped by phosphatidylcholines (PC).^1,2,8^ To characterize the LDs induced by our molecules in Huh-7 cells, we have isolated the LDs using differential centrifugation in a sucrose gradient and the isolated droplets were subjected to liquid chromatography-mass spectral analysis (LC-MS/MS) for specific PCs and TGs.^24,30–33^ PCs possessing carbon chains ranging 34-38 with unsaturation centers 0-4 were noted to be overproduced (∼2-8-fold) than the vehicle control (Figure 2F-H). Interestingly, TGs of 50-carbon chain were enriched, and other examined TGs were lower than DMSO condition (Figure 2F-H). Next, we compared the efficacy of IITK4101 and IITK4102 with OA for promoting lipid droplets in Huh-7 cells under comparable conditions. A striking difference in the overall LDs production was observed between our molecules and OA. While the doses of 2 and 5 µM of OA produced only a minimal or no effect on the number and area of LDs (Figure 2J, J, Extended data figure 1B), at the comparable doses for IITK4101 and IITK4102, a significant increase in the lipid production was recorded (Figure 2I, J, Extended data figure 1B). Together, our screening and subsequent validation has identified small molecules which rapidly induce LDs at sub-micromolar concentrations and are more powerful than fatty acids including OA.

**Figure 2.**
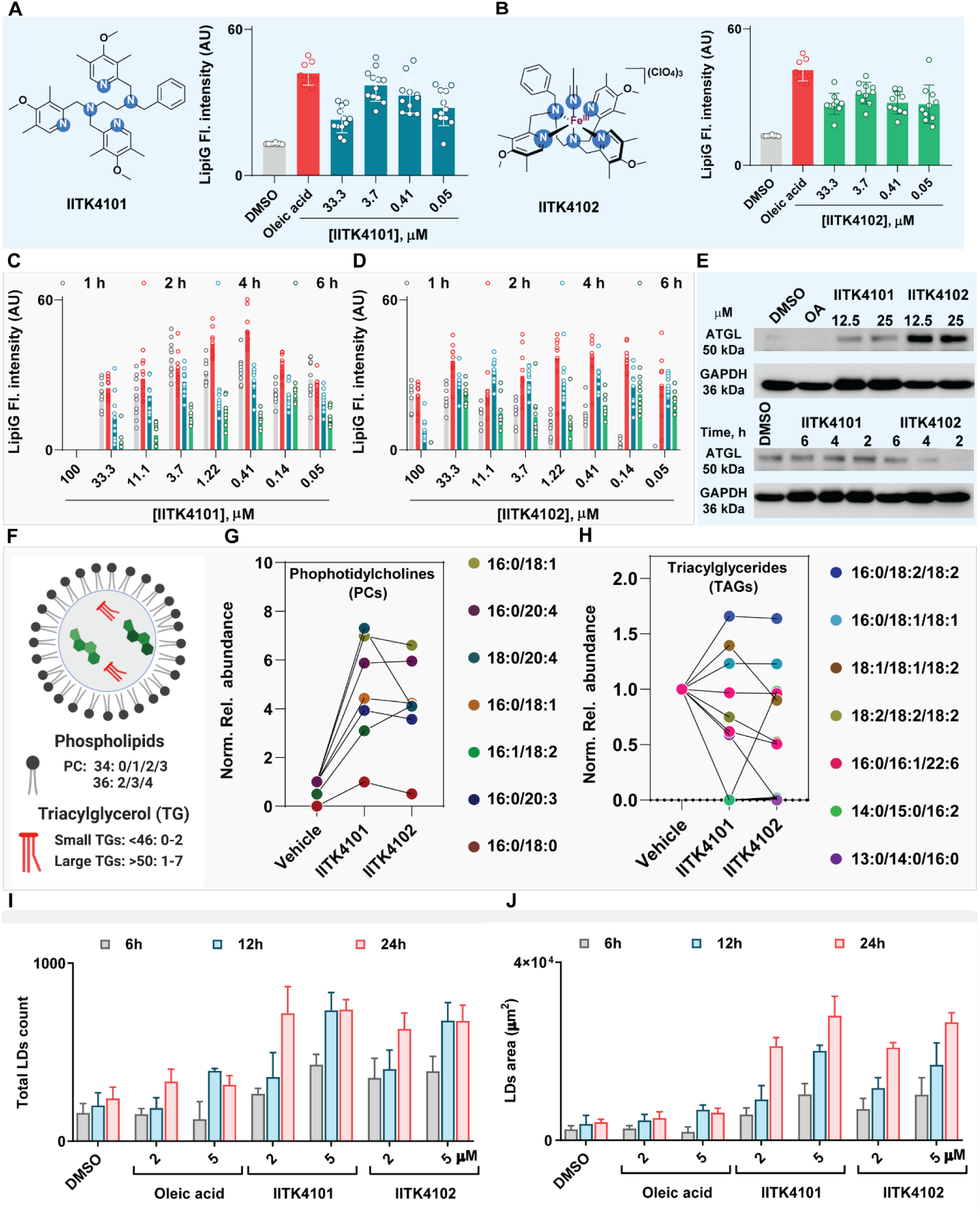
Primary screening hits validation identifies new small molecule inducers of LDs. (A-D) Quantification of LDs using LipiG fluorescence in Huh-7 cells exposed to the lead molecules in dose (measured at 6 h) and time dependent fashion in the indicated concentrations. Oleic acid: 200 µ M. (E). Immunoblotting for ATGL protein in the proteome of Huh-7 cells treated with IITK4101 and IITK4102 in dose (for 6 h) and time-dependent (at 25 µM) fashion. GAPDH is used as a loading control. (F-H) LDs were analyzed using mass spectrometry for selected lipids for the characterization. (I, J). Comparison our lead molecules with oleic acid under comparable concentrations for LDs formation.

### Lipidomics analysis reveals PCs and small TGs enrichment

Next, we extracted the whole lipids and the lipid droplets through a differential centrifugation for lipidomics analysis to investigate the lipid signature.^34^ LC-MS/MS lipidomics analysis of whole lipidome and LDs were performed under comparable conditions. Interestingly in the whole lipidome, enriched levels of PCs were characterized and an overall reduction in the levels of TGs were recorded, globally (Figure 3A, F, G). Conversely, lipidomics analysis of isolated LDs revealed a selective group of PCs being enriched and a majority of the TGs ranging carbon chains 42-54 were globally enriched, which is a hallmark of lipid droplet composition (Figure 3B, C, E-G).^31^ The entire data sets were analyzed for the distribution of various lipids and their compositions, a predominant enrichment of PCs containing lesser number of unsaturation (0-2) were observed (Figure 3C, E-G). On the TGs, a major reduction in the levels of small TGs (C >52) for whole lipidome, and significant enrichment for small TGs in the isolated LDs were recorded (Figure 3B, D, F, G). Overall, the mass spectral lipidomics analysis of composition and lipid signatures of LDs induced by our molecules indicated elevated PCs and small TG levels.

**Figure 3.**
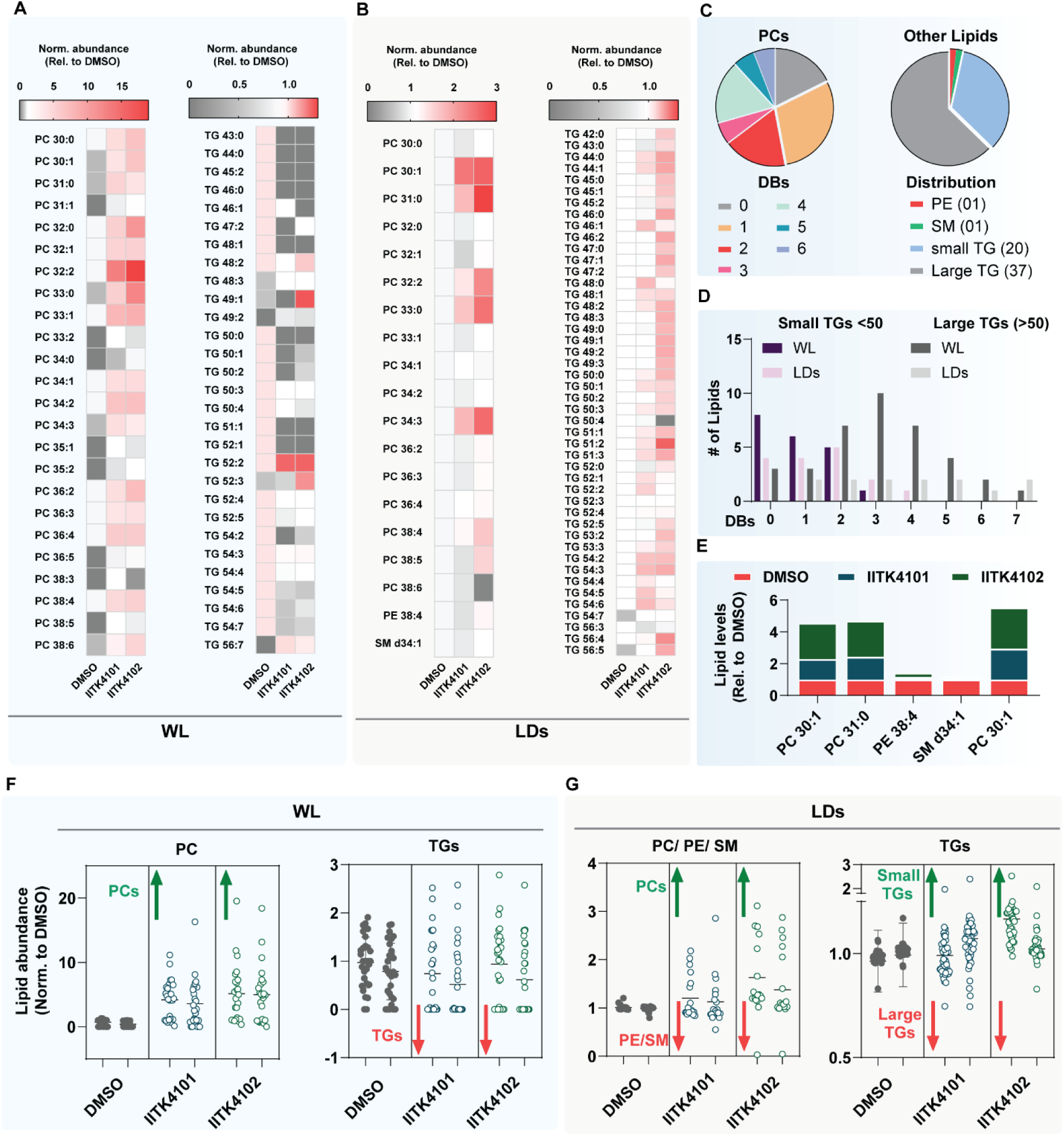
Lipidomics analysis of whole lipidome and isolated LDs reveal upregulation of PCs and small TGs upon small molecules treatment. Lipidome isolated from Huh-7 cells cultured in a 6-well plate treated with IITK4101 and IITK4102 for 6 h at 20 µM for whole lipidome and at 15 µM for isolated LDs. (A, B). Heatmap of lipids abundance obtained from lipidomics (LC-MS/MS) analysis of whole lipidome (A, WL) and isolate LDs (B, LDs). (C, D). Distribution of various class of lipids. PC: phosphotidylcholine; DB: double bond; PE: phosphotidylethanolamine; SM: sphingomyelin; TG: triglycerides (small: < 50 carbon chain; large: >50 carbon chain). (E-G) Plots of entriched lipids in treated conditions compared with DMSO control.

### Our lead molecules in liver and kidney cells promote LDs growth, and disrupted by PC’s addition

In addition to Huh-7, we investigated the potential of our compounds to promote LDs formation in few more cell lines. We selected one more liver cancer cell line (HepG2), kidney cells (HEK293) and osteosarcoma cells for treating with our molecules to study the LDs formation. We have observed a tremendous effect for both IITK4101 and IITK4102 in promoting LDs formation in HepG2 and HEK293 cells to a comparable extent in terms of LDs count, area and size (Figure 4A-D). Interestingly, the treatment of U2OS cells with our molecules had no effect on the LDs formation under comparable conditions (Figure 4A, B). While our molecules are excellent tools for rapid induction of LDs in multiple cells lines at low concentration, which limits the molecules from serving as a universal probe for LDs formation all the cell lines. Lipid droplets are synthesized at the endoplasmic reticulum (ER) and transported to membranes and mitochondria for various functions in cells (Figure 4E, Extended data figure 2A). As shown in the Kennedy pathway, PC synthesized from choline are characterized as a surfactant to prevent LDs coalescence (Figure 4E).^35,36^ To first understand the LDs biogenesis, we assessed the colocalization of LDs with ER in cells treated with our molecules, which showed an excellent overlap, thereby demonstrating an upregulated the LDs biosynthesis driven by our molecules (Figure 4F). Next, the combination of PC along with IITK4101 and IITK4102 during the LDs induction in Huh-7 cells, revealed an overall increase in the count and area of LDs (Figure 4G, H, Extended data figure 2B). However, a sharp decrease in the average size of the LDs were recorded only in the PC-cotreatment condition, which supports the role of PC acting as a surfactant in preventing of LDs coalescence and granularization of existing LDs (Figure 4G, H).^36^ Collectively, IITK4101 and IITK4102 are capable of inducing LDs in liver and kidney cells, and the prevention of LD coalescence with PC serves to characterize LDs.

**Figure 4.**
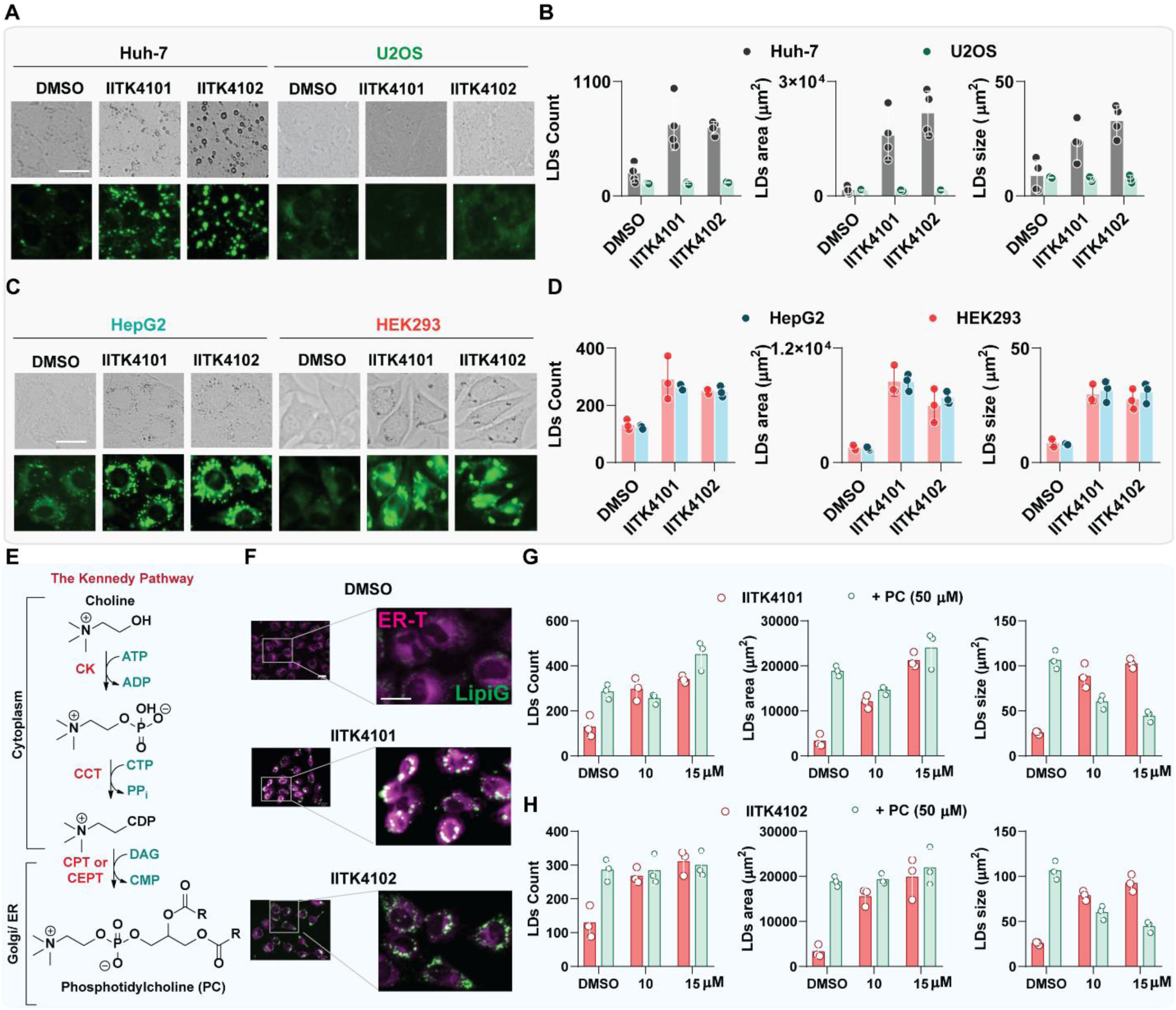
Our chemical tools induce lipid droplets in kidney and liver cells and PC cotreatment prevents the lipid droplet growth. (A, C) Representative images of indicated cell lines visualized for the LDs formation with the compound treatments (15 µM for 3 h), Scale bar: 25 µm. (B, D) Quantification of LDs (count, area and size) induced in Huh-7, U2OS (A-B), HepG2 and HEK293 cells (C-D). (E). Schematic of the Kennedy pathway for the synthesis of PC and relative cellular locations. (F). Colocalization of ER (colored magenta) with LDs (colored green) captured using fluorescence imaging of ER tracker dye and Lipi-green fluorophores, Scale bar: 25 µm. (G, F). LDs quantified after co-treatment of PC with lead molecules to the Huh-7 cells at the indicated concentration for 6 h.

### IITK4101 is maximally effective in LDs induction with iron salt

Given the tremendous effect of IITK4101 and its iron complex, IITK4102 on the LDs formation in Huh-7 cells, we set out to test the effect of other redox and biologically relevant metal ions such as Mn^2+^, Fe^3+^, Cu^2+^, Ni^2+^ and Zn^2+^ chloride salts with IITK4101. When Huh-7 cells are exposed to the indicated metal salts alone, we observed a comparable effect for all the metal salts to that of vehicle control (Figure 5A, B). When these salts are combined with IITK4101, we observed a mild increase in the counts of LDs formation with Mn^2+^, however, the size the area of the lipid droplets were significantly lower than the ligand alone (Figure 5A, B). Combination of Fe^3+^ was the best among the metal salts combined with IITK4101 in all the count, size and area of LDs, albeit marginally lower than the ligand alone (Figure 5A, B) or the complex IITK4102 (Figure 5E, G). In fact, a decrease in the ligand induced LDs count, size and area were recorded for Cu^2+^ or Zn^2+^ salts in combination with IITK4101 (Figure 5A, B). Likely, the complexation with those metal salts is preventing IITK4101 from formation of the active iron-complex with intracellular iron pool. Next, we also assessed a couple more ligands and their iron complexes (IITK4001 to IITK4004), which were previously shown to induce intrinsic apoptosis in cancer cells.^37^ In our assay system, no reliable extent of LDs formation was observed for these compounds, thereby suggesting not any heterocyclic ligands or iron complexes will induce LDs in Huh-7 cells (Figure 5A, B). Further structure activity relationship analysis on IITK4101 and the respective iron complexes could shed more wisdom on the structural handles essential or sensitive for the molecule’s effect on LDs accumulation, which is a bright starting point for future studies. To conclude, IITK4101 exhibited maximal effect on LDs production in Huh-7 cells when combined with iron salt over other biologically relevant metal salts.

**Figure 5.**
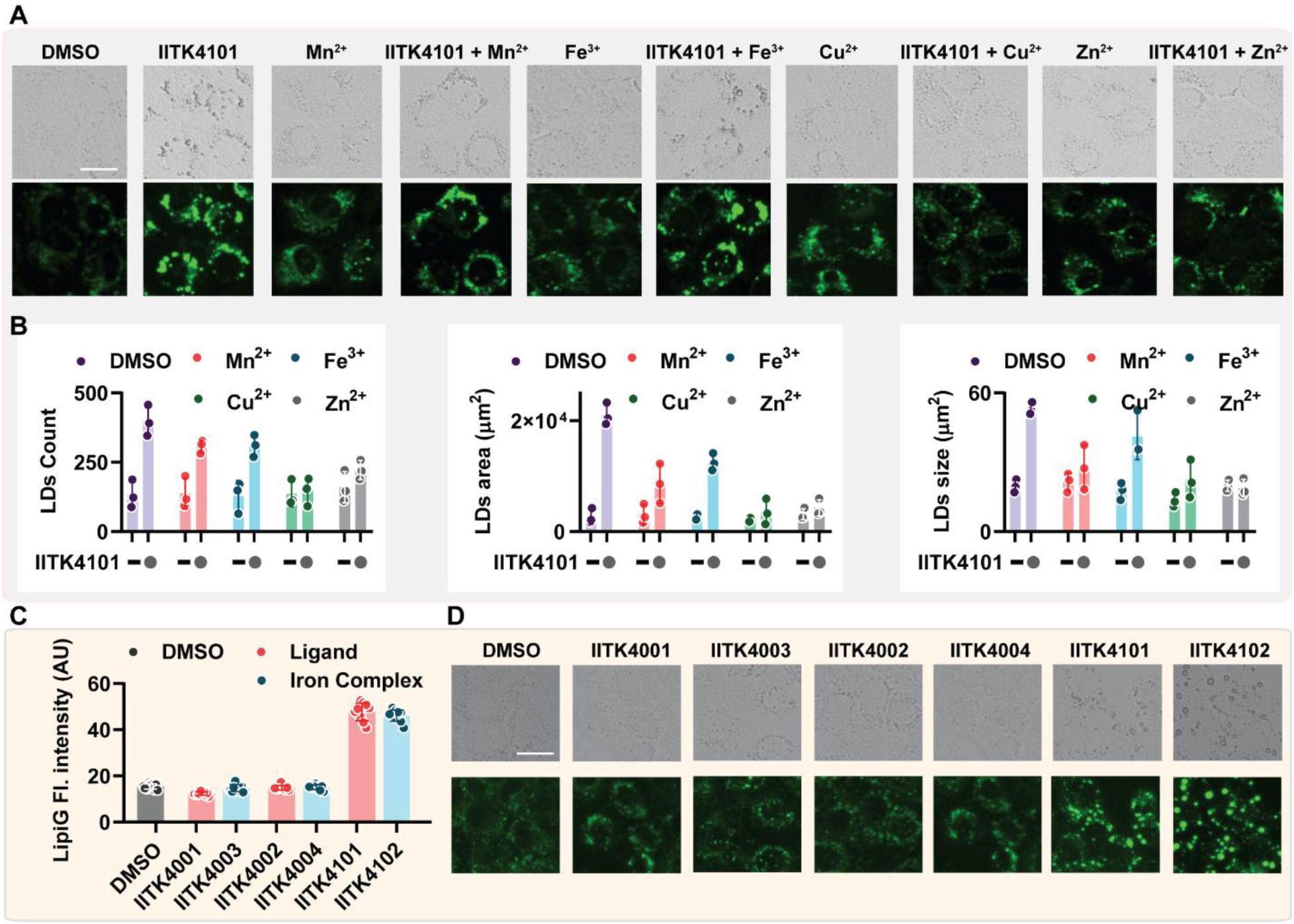
IITK4101 exhibited maximum LDs formation in Huh-7 cells with iron salts. (A, D) Representative images of Huh-7 cells visualized for the LDs formation with the indicated salts (15 µM) and IITK4101 (15 µM), alone or in combination, Scale bar: 25 µm. (B, C) Quantification of LDs (count, area and size) induced in Huh-7 cells after 6 h of compounds’ treatment.

### Mechanism of action studies indicate ROS-mediated lipid peroxidation triggering LDs formation

Iron complexes and the ligands are known to damage mitochondrial function and elevate intracellular ROS.^37^ To test the effect of our molecules on mitochondrial function, mitochondrial membrane potential (MMP) was recorded using a potentiometric fluorescent probe, TMRM (tetramethylrhodamine methyl ester) (Figure 6A). A significant reduction in the overall MMP was recorded in the lead molecule treated conditions and the effect was found to be reversed upon cotreatment with an antioxidant, *N*-acetylcysteine (NAC) (Figure 6A). Subsequent quantification of ATP in cells treated with IITK4101 and IITK4102 revealed a temporary increase in the ATP levels at 30 min and 1 h treatment times (Figure 6B). This could be attributed to the cells attempt at promoting ATP production in order to equilibrate the damaged mitochondrial effect. While continued treatment for 4 h did not affect the ATP levels and a sharp decline in ATP levels was recorded at 6 h treatment (Figure 6B). Next, we measured the extracellular (Figure 6C) and intracellular ROS production (Figure 6D) in the cells treated with IITK4101 and IITK4102 and observed a heavy ROS accumulation in cells, which were found to be suppressed upon the addition of NAC (Figure 6D).^38^ The ROS generated by these molecules were characterized to be less harmful to drive DNA damage in cells (Figure 6E).^39^ Lipid peroxidation induced by intracellular ROS signals the lipids to package themselves into droplets as a protective mechanism. Therefore, we next examined for the lipid peroxidation induced by IITK4101 and IITK4102 in Huh-7 cells (Figure 6F, G). We used TBARS assay which measures the MDA formation as a surrogate for lipid peroxidation and the oxidation of C11-BODIPY, an established lipid peroxidation assay. In both experiments, we observed an enhancement in the levels of oxidized lipids (Figure 6F, G). Lipid droplets generated as an energy source during mitochondrial impairment are known to get associated with mitochondria in an attempt to restore its function.^15,16^ Indeed, tracking mitochondria and LDs using mitotracker and LipiG fluorescent probes in the cells treated with our compounds have shown an excellent colocalization (Figure 6H, Extended data figure 2C). Next, to establish the connection between ROS and LDs formation, we have cotreated the cells with various oxidants and antioxidants along with IITK4101 and IITK4102 and we have recorded a clear trend of increased LD size with oxidants and a reversal in the effect with antioxidants (Figure 6I, Extended data figure 2D). The ensure the contribution arising from lipid peroxidation in LDs generation, we tested RSL3, a ferroptosis inducer functioning via accumulating lipid peroxides,^40^ and we observed the formation of LDs, however lesser extent than out lead molecules (Extended Data Figure 3C, D). Together, the lead molecules promote ROS-mediated mitochondrial impairment to promote LDs formation in cells.

**Figure 6.**
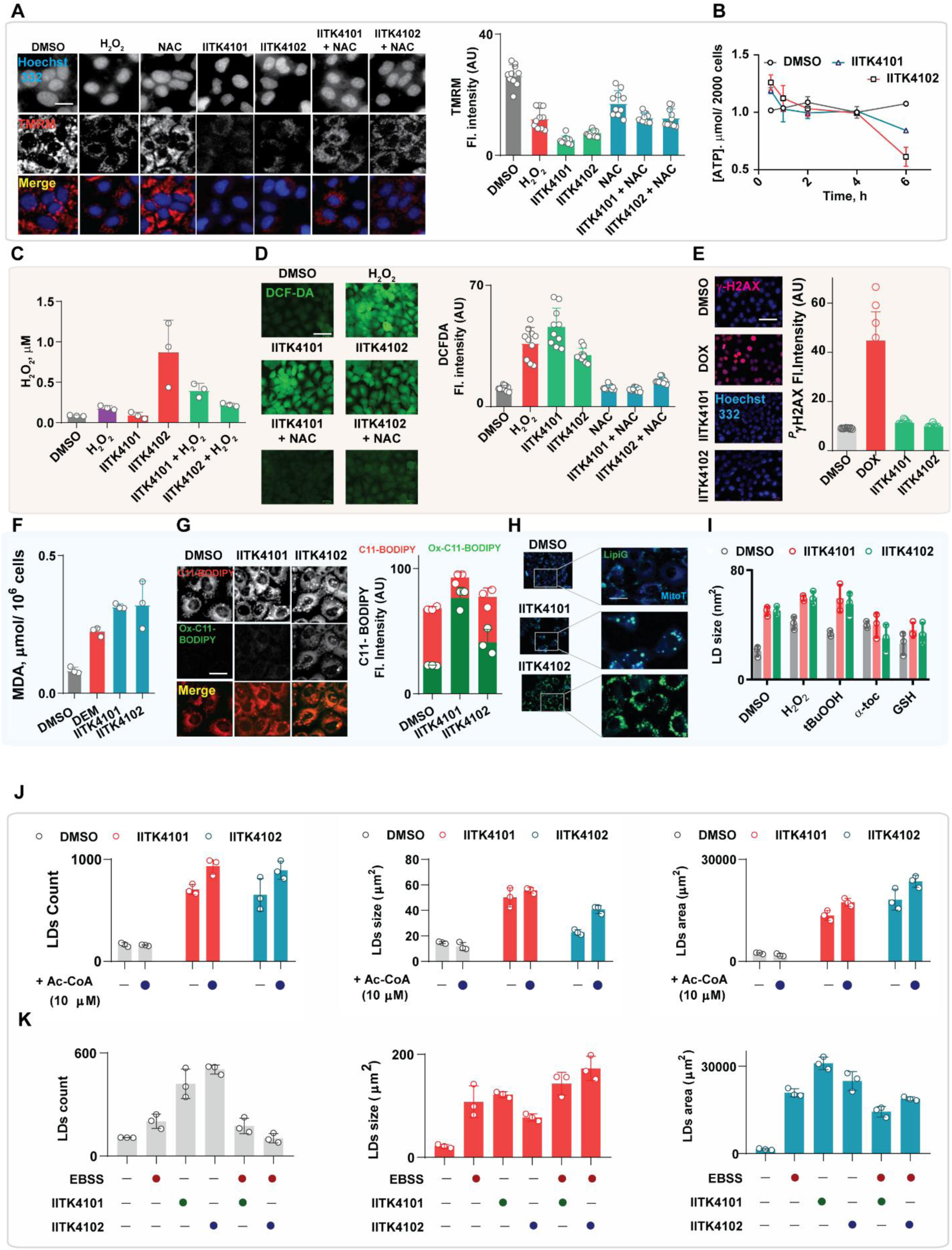
ROS-mediated mitochondria impairment by our lead molecules and lipid peroxidation drives LD formation and starvation prevents the LDs coalescence. (A). Visualization of functional mitochondrial using a potentiometric probe, TMRM and its quantification. Scale bar: 25 µm. (B). ATP quantification after treatment with our lead molecule at indicated time points. (C, D). Quantification of extracellular H_2_O_2_ using Amplex red assay (C), intracellular ROS using H_2_-DCF-DA oxidation assay (Scale bar: 100 µm) and its quantification (D). (E). Assessment of DNA damage response in Huh-7 cells using immunofluorescence imaging for γ-H2AX in Huh-7 cells, Scale bar: 100 µm. (F, G). Lipid peroxidation in Huh-7 cells after treatment with IITK4101 and IITK4102 was assessed using TBARS assay for MDA (malondialdehyde) (F) and C11-BODIPY oxidation (G), Scale bar: 25 µm. (H) Colocalization of LDs on mitochondria in Huh-7 cells captured using mitrotracker (colored blue) and lipigreen fluorescence microscopy, Scale bar: 25 µm. (I - K) Effect of ROS (H), acetyl-CoA (I) and starvation (J) LDs induced by our lead molecules were assessed using combination experiments. In all the experiments presented here, concentrations of compounds used are: IITK4101 & IITK4102: 15 µM, H2O2: 150 µM; α-Toc: 50 µM; *^t^*BuOOH: 30 µM; NAC & GSH: 0.5 mM.

Acetyl-CoA is a source for the fatty acid biosynthesis and its addition is expected to promote the LDs formation further.^41^ Therefore, we tested the effect of Acetyl-CoA on the LDs generated by IITK4101 and IITK4102. With both compounds, an overall increase in the LDs count, size and area are recorded (Figure 6J, Extended data figure 3A). During starvation, the cells themselves drive the LDs formation and transport them to mitochondria for an oxidative metabolism to balance the energy demand.^1,2,12^ In our system, the mitochondrial function is compromised with the compounds treatment, therefore, we expected a further buildup in the size of LDs under starving conditions. To test this, we treated cells cultured in nutrient deprived media (Earle’s balanced salt solution: EBSS) to mimic starvation and treated with IITK4101 and IITK4102. We observed an increase in the number of LDs in nutrient deprived state or in the compounds’ treatment alone (Figure 6K). As expected, a significant decrease in the LDs number and an increase in the LDs size emerged through the fusion of small LDs into large droplets were recorded for our lead molecules only when exposed to starving conditions (Figure 6K, Extended data figure 3B). These studies have clearly demonstrated that both IITK4101 and IITK4102 are inducing LDs through the induction of oxidative insult and the droplets generated could be used by the cells during nutrient deprived conditions.

### Abrupt accumulation of LDs in cancer cell leads to cell death

Cancer cells are sensitive to change in oxidative homeostasis and nutrient stress, due to their high metabolic requirement and proliferation rate.^42^ While LDs accumulation is a hall mark of cancer cells,^43^ we envisioned that the over accumulation of LDs could be lethal to the survival of cancer cells.^44^ Therefore, we treated liver cancer cells, Huh-7 to our lead molecules IITK4101 and IITK4102 for 72 h at indicated concentrations in a dose-dependent fashion (Figure 7A). The cell viability assessment revealed the molecules are exceptionally potent in preventing the growth of cancer cells that exhibited the inhibitory effect at nanomolar concentration (concentration required to prevent 50% of cells’ growth, EC_50_: 410 nM for IITK4101 and 160 nM for IITK4102 against Huh-7 cells) (Figure 7A). We also recorded the molecules toxicity to kidney cells (HEK293) and bone cancer cells (U2OS). Despite a phenomenal induction of LDs in HEK293 cells by our lead molecules, the cells are able to tolerate the LDs accumulation and were found to be highly resistant to our molecules induced toxicity (Figure 7A). This highlights that IITK4101 and IITK4102 are excellent tools for inducing LDs for studying their mechanism of action in cells which are resistant to LDs induced stress. In parallel, we also tested the molecules toxicity to U2OS cells in which the lipid droplet accumulation was not prominent. U2OS cells also observed to exhibit a comparable sensitivity to that of HEK293 that these compounds are not toxic to these cells (Figure 7A). Therefore, LDs accumulation is essential for inducing cytotoxicity in cancer cells by our lead molecules, however, normal cells (kidney cells) capable of tolerating LDs accumulation are not sensitive. Next, we tested the ability of IITK4101 and IITK4102 to prevent cancer cell migration and colony formation in the in vitro conditions. We used scratch assay (wound healing assay) to test the inhibition of cancer cell migration, and we recorded a time and dose-dependent effect of both molecules in preventing the cells movement to that scratch area (Figure 7B, C). Similarly, the clonogenic assay performed to record the inhibition of colony formation in cancer cells also demonstrated a dose dependent effect (Figure 7D, E). Next, to verify the contribution of oxidative stress in promoting cell death, we performed cell viability experiment of our lead compounds (IITK4101 and IITK4102) with various oxidants and antioxidants (Figure 7F). Other cell death mechanisms such as apoptosis and necroptosis were eliminated after observing no significant cell death prevention recorded when IITK4101 and IITK4102 were independently combined with QVD-OPh, a pan caspase inhibitor for apoptosis) and necrrostain-1 (RIPK1 inhibitor for necroptosis) (Figure 7F). Iron mediated accumulation lipid peroxidation is well characterized to promote ferroptotic cell death. ^40^ Therefore, it is reasonable to expect our lead molecules to induce ferroptosis for preventing cancer cell growth. However, a cotreatment of our compounds with ferroptosis modulators (erastin, RSL3 and FIN56 as ferroptosis inducers and ferrostatin-1 and liproxstatin-1 as ferroptosis inhibitors) showed no alteration in the overall cell death profiles induced by IITK4101 and IITK4102 (Figure 7F). Here, we demonstrate that rapid and heavy accumulation of LDs could be used as a potential therapeutic strategy to prevent cancer growth and the mechanism of actions for the lead molecules stand not related to oxidative stress induction.

**Figure 7.**
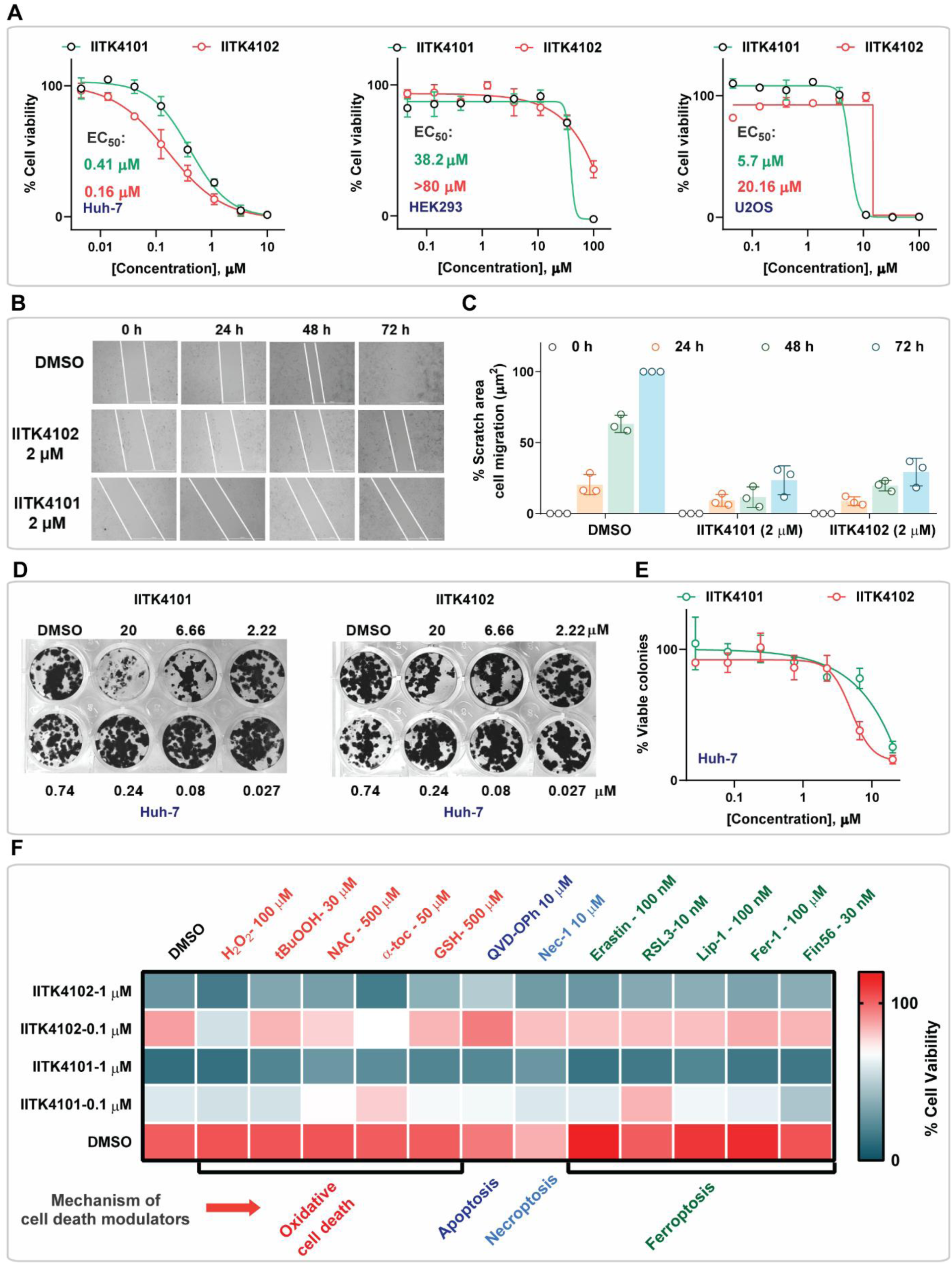
LDs inducers were potent inhibitors of liver cancer cell proliferation. (A). Cell viability plot of Huh-7 cells after treatment with our lead molecules in an 8-point-dose for 72 h, the data is normalized to DMSO. (B-E). Influence of IITK4101 and IITK4102 on cell migration (scratch assay, B, C) and colonization of Huh-7 cells (clonogenic assay for 72 h, D, E) was assessed after indicated time points. (F). Combination cell viability data recorded for Huh-7 cells after 72 h of treatment with IITK4101 and IITK4102 in combination with various cell death pathway modulators at indicated concentrations.

### Lead molecules promote lysophagy in cancer cells

Autophagy is a programmed mechanism that clears damaged proteins, organelle and cells, which is frequently encountered in biological systems associated with lipid droplets.^45,46^ Therefore, we suspected that our LDs inducing small molecules, IITK4101 and IITK4102 could promote hyperactive autophagosome formation to induce cancer cell death. Lipidation of LC3B-I protein to form LC3B-II is one of the canonical biomarkers for the characterization of autophagy activation.^47,48^ Along with our compounds, we chose to induce the nutrient deprived state as a positive control for LC3B-II formation, and the autophagosome-lysosome fusion inhibitor, bafilomycin A1 (Baf) was used as another control for enriching lapidated LC3B-II.^49,50^ Immunoblotting for LC3B-I and lipidated LC3B-II in Huh-7 cells proteome after treatment with chosen conditions for 2 h, the positive controls worked as expected, however, no effect was recorded for IITK4101 and IITK4102 (Figure 8A). Interestingly, even the Baf and EBSS cotreatment with our compounds also did not show any lipidated LC3B-II presence in the proteome (Figure 8A). After 2 h of exposure of Huh-7 cells to our compounds, we recorded a significant amount of LDs formation, however, no symptom of LC3B-II production has led us to conclude the induction of autophagy is an unlikely mechanism. Then, we prolonged the exposure to 4 h and observed a sharp increase in the LC3B-II levels, and the cotreatment with EBSS or Baf has produced higher levels of lipidated LC3B-II (Figure 8A). This could be due to a delayed on-set of autophagy activation or may be a causal effect of huge stress induced by abrupt accumulation of LDs. To clarify this, next we focused our attention on TFEB protein (Transcription factor EB) which gets translocated into the nucleus during stress conditions, particularly in nutrient deprived state, and activates genes responsible for lysosome and autophagosome biogenesis.^51–53^ We used LLOMe (l-leucyl-l-leucine methyl ester) as a positive control, a known inducer of TFEB translocation through lysosomal damage^54^ and rapamycin (Rap, autophagy inducer) was used as a negative control (Figure 8B, C, Extended data figure 4A). Here, we observed a sharp increase in TFEB translocation than DMSO in all conditions supporting lysosomal damage and activation of autophagy in cells (Figure 8B, C). However, both autophagy inducer (Baf) and inhibitor (Rap) had no effect on the antiproliferative activity of IITK4101 and IITK4102. Collectively, we have concluded autophagy is an unlikely mechanism of action for our lead molecules in inducing cancer cell death.

**Figure 8.**
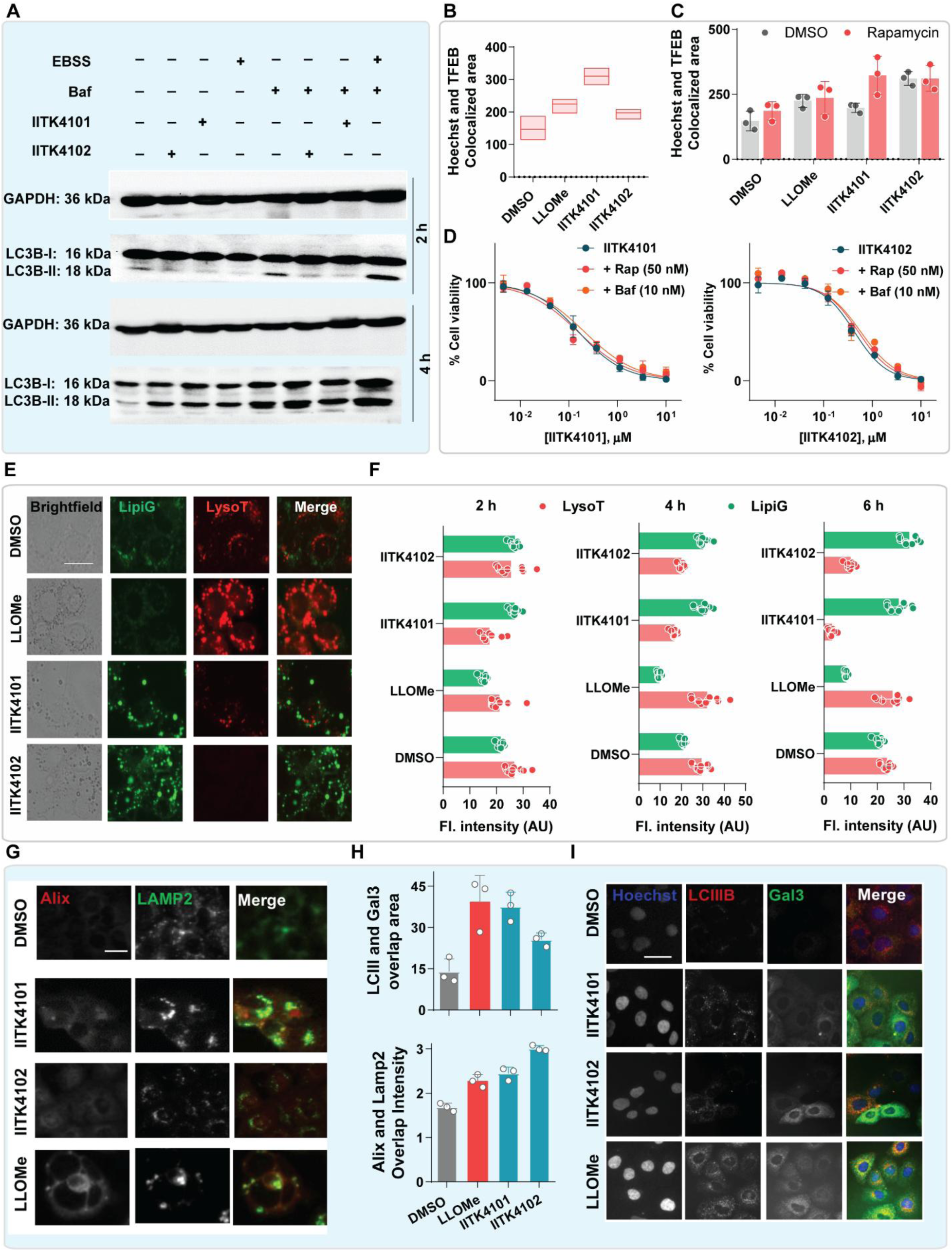
LDs inducing molecules promote lysophagy in Huh-7 cells. (A). Immunoblotting for LC3B-I and LC3B-II in huh-7 cells proteome collected after treatment with IITK4101 and IITK4102 under various combinations. starvation: EBBS medium; Baf: bafilomycin A1. GAPDH is used as control. (B, C). Translocation of TFEB from cytoplasm to the nucleus was quantified using Hoechst 332 colocalization area upon of Huh-7 cells treatment with IITK4101 and IITK4102 (15 µM) alone (B) and in combination with rapamycin (Rap, 50 nM) (C). LLOMe (500 nM) was used as positive control. (D). Cell viability dose-curve of Huh-7 cells after treatment with IITK4101 and IITK4102 alone and in combination with Rap and Baf for 72 h, the data is normalized to DMSO. (E, F). Effect of LDs on lysosomes was assessed in Huh-7 cells after 6 h treatment with IITK4101 and IITK4102 (15 µM) using fluorescence imaging for LDs (LipiG) and lysosomes (Lysotracker; Scale bar: 25 µm) and its quantification at 2, 4 and 6 h treatment conditions (F). (G-I). Colocalization of lysosomal repair proteins Gla3 and Alix with Lamp2 and LCIII in Huh-7 cells for the indicated conditions were assessed using immunofluorescence imaging, Scale bar: 25 µm.

Our observation of TFEB translocation to activate lysosome biogenesis in the compound treatment conditions prompted us to investigate lysosomes in cells health. Aggregation of α-synuclein protein were known to damage lysosomes and induce lysosomal death, a mechanism called lysophagy.^55^ Therefore, we suspected that the aggregation of LDs could potentially induce lysophagy in our system. First, we assessed the colocalization of lysosomes with LDs using lysotracker probe and LipiG and we observed an exceptional colocalization (Figure 8E, F, Extended data figure 4B-D). Interestingly, the treatment of LLOMe leading to lysosomal damage has activated the lysosomal biogenesis in cells which was evident from overall increase in the staining for lysosomes (Figure 8E, F, Extended data figure 4B-D). However, in our compound treatment conditions, we observed a drastic decrease in the staining for lysosomes highlighting the loss of active lysosomes in a time-dependent fashion (Figure 8E, F, Extended data figure 4B-D). The loss of lysosomes was directly correlated with the extent of LDs production in the cells (Figure 8E, F, Extended data figure 4B-D) and we observed no recovery in the lysosome biogenesis on prolonged exposure. Upon lysosomal damage, lysosome repair proteins will be activated by TFEB and recruited to the damaged lysosome for repair.^56–61^ The endosomal sorting complex required for transport (ESCRT) is recruited to the damaged lysosomal membrane for a repair and to establish the lysosomes quality control.^58–61^ We chose to observe ESCRT recruiting proteins galectin3 (Gal3) and ALIX colocalization to the lysosomes to observe the lysosomal repair mechanism activation.^56,61,62^ Immunofluorescence imaging of cells treated with LLOMe as a positive control exhibited to sharp increase in the Gal3 and ALIX levels and their colocalization at the lysosomes and validated the assay system (Figure 8G-I). Under comparable conditions, IITK4101 and IITK4102 were equally effective in recruiting Gal3 and ALIX to the lysosomes, thereby establishing lysosomal damage in the compound treated conditions (Figure 8G-I). These experiments have prominently established the cancer cell death mechanism induced by IITK4101 and IITK4102 is lysophagy.

## Discussion

LDs are implicated to play a major part in several health and disease conditions, including cancer, diabetes and obesity.^63–65^ While lipid droplets existence is proposed to be prevalent in majority of cell types to tackle metabolic requirements, the precise role and importance of LDs remains elusive.^1,2,63^ Chemical tools that are capable of rapidly inducing LDs at lower doses could facilitate the understanding of LDs in disease and health. Therefore, we developed a high-content imaging-based phenotypic screening for the identification of structurally new small molecules for LDs induction in Huh-7 cells. We used a synthetic fluorophore, lipi green for a selective labelling of lipid droplets and for reliably quantifying the extent of LDs generated in cells (Figure 1A, B).^27^ We aimed to take advantage of the oxidative stress mediated lipid peroxidation as a signaling cue for promoting LDs formation in cells. Our screening of RSL3, a lipid peroxidation inducing molecule that triggers ferroptosis in cancer cells,^40^ was found to promote LDs formation in Huh-7 cells (Extended data figure). We reported that *N*-heterocyclic small molecules capable of forming metal complexes with iron elevated oxidative stress in cells.^37^ Using our screening platform, we screened *N*-heterocyclic ligands spanning bidentate, tridentate and tetradentate coordination sites with metals. Our screening and spatiotemporal analysis has identified small molecules functioning at less than a micromolar dose and within 2 h of treatment (Figure 2A-J). A fluorescence imaging-based assay platform could be used for the identification of new LDs inducers in cells.

Traditionally, OA at high micromolar doses (∼200 µM) is used as LD inducing fatty acid in various cell-based models and in animal experiments. Any knowledge on optimal number of unsaturation sites in fatty acids for effectively inducing LDs is not available. Using our screening platform, we have screened fatty acids possessing various number of double bonds, which included stearic acid, oleic acid, linoleic acid, linolenic acid and arachidonic acid possessing 0, 1, 2, 3 and 4 DBs, respectively. From the complete analysis of LDs quantification data in Huh-7 cells, system containing 2-DBs, linoleic acid (LOA) was identified to be the best in rapidly inducing LDs over other fatty acids, including OA under comparable conditions. LOA was also found to be least toxic to cells even at the highest concentration tested (200 µM) over other higher polyunsaturated fatty acids (Figure 1C, D).

The versatility of our lead molecules inducing LDs in kidney cells and liver cells highlights the potential of these compounds to be employed in studying LDs effect in specific disease models in different cell types. While the inability of these molecules in promoting LDs formation in osteosarcoma is a limitation, a universal application of these compounds as LDs inducers needs to be investigated in more cell types with additional molecules. The mechanism of action of these molecules for triggering LDs formation was linked to the mitochondrial impairment mediated lipid peroxidation (Figure 6A-B). A natural extension that the cells resistant to oxidative stress are unlikely to respond to these molecules for promoting LDs formation, which remains to be studied. The mechanism of stress induced LDs formation is established for our compounds when combined with acetyl CoA and nutrient deprived conditions using EBSS medium (Figure 6J, K). The induction of cancer cell death when exposed to our lead molecules in which abrupt elevation of LDs formation is recorded, is not surprising. However, despite the heavy accumulation of LDs in human embryonic kidney cells, they were extremely resistant to our compounds’ induced lipid stress and toxicity. This exciting finding highlights the possibility of inducing LDs in cells without compromising on cell health to study the role and impact of LDs accumulation on health and disease.

Cancer cells undergo severe stress and alteration in the metabolic network, which is highly complex and sensitive to change in the stress conditions.^23,43,44^ We demonstrated that the induction of LDs in cancer cells by our molecules prevented their proliferation and addition of neither oxidants nor antioxidants had any effect on the small molecules’ toxicity (Figure 7F). Therefore, we concluded the toxicity is unlikely to be due to the mild oxidative stress induced by the molecules. This further supported the weak toxicity profile observed for IITK4101 and IITK4102 in HEK293 cells, which is more resistant to relatively mild change in oxidative stress than cancer cell lines (Figure 7A). Interestingly, these compounds were highly effective in preventing the cancer cell movements and colonization in addition to their antiproliferative activity (Figure 7B-E). Unless the cells are sensitive to rapid accumulation of LDs, their growth is unlikely to be disturbed by our molecules.

Finally, we demonstrated the mechanism responsible for the cancer cell death induced by compounds was due to activation of lysophagy upon the heavy production of LDs in cells. Induction of autophagy in LDs enriched condition is well documents, however, in our case, a sharp decrease in the lysosomal count was recorded when exposed to our compounds. This is not entirely surprising since, under a similar circumstance of α-synuclein aggregation in cells led to lysophagy.^55^ This observation was supported by the recruitment of lysosomal repair proteins, ALIX and Gal3 to the lysosomes during the compound treatment (Figure 8G-I). These results provide new small molecule tools for reliably inducing lipid droplets in cells for studying their downstream actions. We demonstrated the induction of lipid droplet accumulation using small molecules in liver cancer cells (Huh-7) could trigger lysophagy to prevent their growth, migration and colonization.

## Methods

### Statistics and reproducibility

Data were expressed as mean ± s.d. and P values were calculated using an unpaired two-tailed Student’s t-test for pairwise comparison of variables with a 95% confidence interval and n − 2 degrees of freedom, where n was the total number of samples, in all Figures. All the biological studies were performed in Huh-7 cell line and were representative of two or more independent experiments with a minimum of triple technical replicates.

### Cell culture

Huh-7, U2OS, HepG2, and HEK293 cells were grown in DMEM (Dulbecco’s Modified Eagle Medium Gibco). DMEM was supplemented with 10% FBS (Foetal Bovine Serum-Gibco) and 1% PenStrep (Pencillin Streptomycin-Gibco). Cells were incubated at 37 °C, 5% CO_2_ (Forma-steri-cycle CO_2_ incubator - ThermoFisher Scientific) for growth/treatment.

### Dilution procedure of metal complexes/ligands/test compounds

The percent stock concentration of all the molecules was 100 mM and made in 100% Dimethyl sulfoxide (DMSO)(SRL). The parent stock was stored in −20 °C, the sub stocks and further dilutions were always freshly made in DMEM media just before the experiment

### General protocol of staining the lipid droplets with lipi green

In 96 well plates (NEST), 20000 Huh-7 cells/well were seeded in 200 µL DMEM media and incubated for 24 h. Then the cells were treated with test compounds with the mentioned concentrations and incubated for various time periods. Then the cells were stained with lipi green (1 µM) and further incubated for 30 minutes. Post incubation, media was removed, cells were washed with 1x PBS (100 µL) for twice and the images were captured using fluorescent microscope (Bio-Rad ZOE fluorescent cell imager) in the green channel (λ_ex_ 480 nm; λ_em_ 517 nm). The fluorescence intensity quantification of lipi green, total count of lipid droplets, area and the size of lipid droplets in the cells were calculated by using ImageJ software and the graphs were plotted by using Graphpad Prism 9.0 software. The data provided were representative of three or four independent experiments.

### Concentration of compounds used for staining lipid droplets

For screening studies, Huh-7/ HEK293/ U2OS/ HepG2 cells were treated with 10 µM of heterocyclic compounds, IITK4101, IITK4102, IITK4001, IITK4002, IITK4003, IITK4004, 200 µM of stearic acid, oleic acid, linolenic acid, and 100 µM of cholesterol (SRL) with 0.01% DMSO as negative control. Cells were incubated for 3 hours for treatment and taken forward for lipi green staining. For the preliminary dose and time dependent studies, 33.3 µM to 0.05 µM (3-fold dilutions) of IITK4101, IITK4102 with 200 µM of oleic acid as positive control and 0.02% DMSO as negative control were taken. The cells were incubated for 3 h with test compounds, and the lipid droplets were stained with lipi green. Further, dose-dependent studies were carried out with eight-point-three-fold dilutions with concentrations ranging from 200 µM to 0.27 µM of stearic acid, oleic acid, linoleic acid (BLD Pharma), linolenic acid, arachidonic acid (BLD Pharma), cholesterol and 100 µM to 0.05 µM of IITK4101/IITK4102.with 0.02% DMSO as negative control. Huh-7 cells were incubated with fatty acids for 6 h, 12 h and 24 h. With IITK4101/IITK4102, cells were incubated for 1 h, 2 h, 4 h, 6 h. For the combination treatment studies, 15 µM of MnCl_2_, FeCl_3_, CuCl_2_, ZnCl_2_, IITK4101, IITK4102, 10 µM of Acetyl CoA (Sigma), 50 µL of EBSS (Gibco)+ 5 µM of IITK4101/IITK4102, H_2_O_2_-150 µM (TCI), ^t^BuOOH-30 µM (GLR), *α*-tocopherol-50 µM (SRL), Glutathione-500 µM (SRL), Phosphatidylcholine-50 µM (SRL) and were used. In comparison studies, 5 µM and 2 µM of oleic acid, IITK4101/IITK4102, 50 µM of stearic acid, oleic acid, linoleic acid, linolenic acid, arachidonic acid and cholesterol were used. In all the comparison and combination lipi green staining studies, 0.01% DMSO was used as negative control, and the cells were incubated for 3 h except 6 h in case of EBSS.

### Isolation of lipid droplets

In T-25 flask (Thermo), 1.5 million Huh-7 cells were plated and incubated for 24 h. They were treated with 15 µM of IITK4101/IITK4102, 100 µM of oleic acid separately for 6 h and scrapped with 1 mL of ice cold 1x PBS. The cell suspension was centrifuged at 4 °C for 10 min and the supernatant was discarded. The pellet was resuspended in 1 mL of HLM buffer (20 mM of Tris-HCl (SRL) at pH-7.4, 1 mM of EDTA (SRL) at pH-8 in milli-Q-water) with 1x PPI (Sigma) and incubated on ice for 10 min. Then the cells were probe sonicated for 1 minute at 5% power rate and centrifuged for 10 minutes at 4 °C to collect the post nuclear supernatant. To the supernatant, 200 µL of 60% sucrose (sigma) (dissolved in HLM buffer) was added and resuspended thoroughly. Then a layer of 300 µL of 5% sucrose solution (made with HLM buffer) was added on top of 60% sucrose undisturbed. This was further covered with 300 µL of HLM buffer. Then the centrifugation was performed at 28,000g for 30 min at 4°C. The upper layer which contains lipid droplets was carefully collected and the centrifugation step was repeated by the addition of 300 µL of HLM buffer to maximize the lipid droplet collection. Lipid droplets from HLM buffer were extracted by adding 600 µL of chloroform (Rankem) and 400 µL of methanol (Rankem), vortexed briefly and centrifuged at 2800xg for 5 min at 4°C. The organic phase was collected, and the same was repeated by adding 600 µL of chloroform and centrifuged further. The final lipid droplets samples were dried under N_2_ gas and taken forward for lipidomics.

### Isolation of whole lipidome

In 6-well plate (NEST), 1 million Huh-7 cells/well were seeded and incubated for 24 hours. They were treated with 20 µM of IITK4101/IITK4102 separately for 6 h and scrapped with 1 mL of ice cold 1x PBS separately. The cell suspensions were added with 2 mL of chloroform and 1 mL of methanol, vortexed and centrifuged at 2800xg for 5 min at 4°C. The organic phase was collected, 50 µL of formic acid was added and the centrifugation step was repeated by adding 4 mL of chloroform. The organic phase was again collected and was dried under N_2_ gas. The dried lipids were solubilized by adding 2:1 (v/v) chloroform and methanol for lipidomics analysis.

### Lipidomics analysis

To study the lipid profile of our samples (whole lipidome and lipid droplets), an untargeted lipidomics was performed. An Agilent 1290 Infinity II Ultra-High-Performance Liquid-Chromatography (UHPLC) system connected to an Agilent 6546 quadrupole time-of-flight (QTOF) mass spectrometer was used as the analytical platform for data collection. Extracted samples in methanol: isopropanol (1:1, v/v) were placed in an autosampler (maintained at 4 °C) attached to the Agilent 1290 Infinity II system. The samples were injected into the LC-MS/MS system (5 µL) and separated the lipids using Agilent ZORBAX Eclipse Plus-C18 (2.1 × 50 mm, 1.8 Micron) column maintained at 60°C (column oven temperature). The mobile phases used for the separation consisted of (A) 10 mM ammonium formate (SRL) in 9:1 water/MeOH (v/v) and (B) 10 mM ammonium formate, in 2:3:5 acetonitrile/MeOH/isopropanol (v/v). The elution gradient started at 70% of B at 0–2 min, 82% at 2–15 min, 90% at 15–28 min. 98% at 28–34 min, 100% at 34–43 min. Starting conditions were recovered at minute 43, followed by a 5 min re-equilibration time, a total running time of 47.1 min. The flow rate during the analysis was kept constant at 0.3 mL/min.

The parameters of an Agilent 6546 QTOF mass spectrometer with a dual AJS ESI ion source were: 750 V octopole radio frequency voltage, 3500 V capillary voltage, 150 V fragmentor voltage, 65 V skimmer, 10 L/min nebulizer gas flow, 200 °C gas temperature, 35 psi nebulizer gas pressure, 12 L/min sheath gas flow, and 300 °C sheath gas temperature. Auto MS/MS mode was used to do the analysis of the samples. MS acquisition in positive ESI mode conducted in full scan mode from 100 to 1200 m/z at a scan rate of 3 spectra/s. In the positive ion mode, MS/MS acquisition was carried out between 80 and 1200 m/z at a scan rate of 4 spectra/s, with collision energies of 20 eV and 35 eV. The software selects the six more intense precursor ions for each measurement, which were fragmented to obtain the spectrum for a specific time. The raw data from the mass spectrometer were analyzed by using Lipid annotator software from Agilent technology. Note: All samples were injected in technical duplicate into the LC-MS/MS.

### Mitochondrial membrane potential analysis

For mitochondrial membrane potential analysis, 25000 cells per well were plated in clear flat bottom 96 well plates (NEST) in 200 µL DMEM and incubated for 24 h. After incubation, cells were treated for 90 minutes with 0.02% DMSO, IITK4101-15 µM, IITK4102-15 µM, H_2_O_2_ - 150 µM, NAC-500 µM (SRL), IITK4101 + NAC, IITK4102 + NAC. Post treatment, wells were washed once with 100 µL 1x PBS. Cells were co-stained 30 nM of Tetramethyl rhodamine, methyl ester percholate (TMRM) - (TCI) and 1µg/mL Hoechst 33342 (Thermo) in 50 µL DMEM for 30 minutes in incubator. After staining, cells were washed with 100 µL 1X PBS for twice. and the images were captured using fluorescent microscope e (Bio-Rad ZOE fluorescent cell imager) red channel (λ_ex_-556 nm; λ_em_-615 nm) for TMRM, blue channel (λ_ex_-355 nm; λ_em_-433 nm) for Hoechst 342. Images colocalizations and fluorescence intensity quantifications were performed by using ImageJ software. Bar graphs were plotted by using Graphpad Prism 9.0 software. The data provided were representative of three or four independent experiments.

### Quantification of cellular ATP

For the quantification of Adenosine-5’ triphosphate in Huh-7, ATP magnesium salt (sigma) standard curve generation experiment was performed. 8 different concentrations of ATP were prepared in 50 µL 1X PBS in triplicates as follows, 100 μM, 25 μM, 6.25 μM, 1.56 μM, 0.39 μM, 0.097 μM, 0.024 μM, 0.006 μM, 0 μM. 25 µL of 3X CellTiter-Glo reagent was added to all these ATP samples (final concentration of CTG was1X). The luminescence readings were recorded in multimode plate reader. Background luminescence (0 μM ATP with only 1X CTG) was subtracted from all the other samples. XY scatter graph was plotted in microscoft excel by taking concentrations of ATP on X-axis and luminescence readings on Y-axis. Straight line equation, Y=mX+C was obtained from the graph. For measurement of total intracellular ATP levels in cells, 2000 cells per well in 200 µL DMEM were plated in white flat bottom 96 well plates and incubated for 24 h. After incubation, cells were treated with 1 µM, 5 µM, 10 µM of IITK4101/IITK4102 for 2 h, 4 h and 6 h. After treatment, the media was removed from wells, 75 µL of 1X CellTiter-Glo reagent was added to each well incubated at room temperature for 10 minutes and luminescence readings were taken by using multimode plate reader. Background luminescence was removed and the concentrations of ATP in Huh-7 cells which were treated with various concentrations of compounds were calculated by using Y=mX+C equation of ATP standard curve, where Y was luminescence reading, m was slope, X was concentration of unknown ATP and C was constant. The data provided were representative of three or four independent experiments.

### Intracellular quantification of ROS by DCFDA

For measurement of total intracellular ROS levels in Huh-7 cells, 20000 cells per well were plated in clear flat bottom 96 well plates in 200 µL DMEM and incubated for 24 h. After incubation, cells were treated for 90 minutes 0.02% DMSO, H_2_O_2_ - 150 µM, NAC - 500 µM, IITK4101/IITK4102 - 15 µM, IITK4101+NAC, IITK4102+NAC. After treatment, cells were stained with 2 µM (final concentration) of 2’,7’-Dichlorofluorescein 3’,6’-diacetate (DCFDA) (Thermo) (dissolved in DMSO) was directly added to cells 200 µL of DMEM media for 5 minutes.in the incubator. After staining, cells were washed twice with 100 µL of 1X PBS, cells were observed under microscope and images were captured using fluorescent microscope (Bio-Rad ZOE fluorescent cell imager) in the green channel (λ_ex_ 480 nm; λ_em_ 517 nm). Fluorescence intensities were quantified by using ImageJ software. Bar graphs were plotted by using the Graphpad Prism 9.0 software. The data provided were representative of three or four independent experiments. The data provided were representative of three or four independent experiments.

### Pγ-H2AX immunofluorescence imaging

For Pγ-H2AX immunostaining experiment in Huh-7 cells, 20000 cells per well were plated in clear bottom 96 well plate and incubated for 24 h. They were treated in triplicate with DMSO, 10 μM of IITK4101/IITK4102, 1 μM of Doxorubicin for 2 hours in the incubator. Post treatment, cells were washed with 200 μL of ice cold 1x PBS and fixed with 100 μL of 4% formaldehyde for 15 minutes at room temperature. Then the cells were washed with 200 μL of ice cold 1x PBS for twice. Then 100 μL of ice cold 100% methanol was added to the cells and incubated at −20 °C for 10 minutes. They were again washed with 200 μL of ice cold 1x PBS for thrice. Pγ-H2AX (Cell Signalling Technology) was diluted in 1:50 ratio by using antibody dilution buffer (1% BSA in 1X PBS) and 50 μL of diluted antibody was added to the cells and incubated for 1 hour in dark at room temperature. In the last 10 minutes, the cells were stained with 1 μg/mL Hoechst 342 stain. Post staining, cells were washed thrice with ice cold 1X PBS and images were captured under fluorescent microscope (Bio-Rad ZOE fluorescent cell imager) red channel (λ_ex_ −556 nm; λ_em_ −615 nm) for TMRM, blue channel (λ_ex_ −355 nm; λ_em_ −433 nm) for Hoechst 342. Images colocalizations and fluorescence intensity quantifications were performed using ImageJ software. Bar graphs were plotted by using the Graphpad Prism 9.0 software. The data provided were representative of three or four independent experiments. The data provided were representative of three or four independent experiments.

### TBARS-lipid peroxidation assay

For lipid peroxidation assay in Huh-7 cells, 0.35 million cells per well were plated in clear bottom 12 well plate (NEST) and incubated for 24 hours. They were treated in triplicate with DMSO, 20 μM of IITK4101/IITK4102, 4 mM of Diethyl maleate for 6 hours in the incubator. Post treatment, cells were washed with ice cold 1X PBS. They were scrapped by adding 200 μL of 2.5% trichloroacetic acid (SRL) and centrifuged for 15 minutes at 13,000g. The supernatant was collected and 200 μL of 15% TCA and 500 μL of 0.67% thiobarbituric acid (TBA) (GLR) were added to the same. They were vortexed were heated at 95 °C for 30 minutes and allowed to cool down to room temperature. To all the samples, 500 μL of n-butanol (Rankem) was added and let the aqueous and organic phases got separated. 200 μL of organic phase of all the samples were added to various wells of clear bottom 96-well plate and fluorescence readings were recorded at λ_ex_ −530 nm and λ_em_ −550 nm in multimode plate reader. For the quantification of MDA, malondialdehyde tetrabutyl ammonium salt (sigma) standard curve generation experiment was performed. 5 different concentrations of MDA were prepared in 2.5% TCA in triplicates as follows, 2.5 μM, 1.25 μM, 0.625 μM, 0.312 μM, 0 μM. To these, 100 μL of 0.67% TBA was added and heated at 95 °C for 30 minutes. The samples were allowed to cool down and fluorescence readings were recorded as described above. Concentration of MDA in the samples was calculated by using Y=mX+C equation of MDA standard curve, where Y was fluorescence reading, m was slope, X was concentration of unknown MDA and C was constant. The graphs were plotted using Graphpad Prism 9.0 software. The data provided were representative of three or four independent experiments.

### C11-BODIPY staining

For the measurement of lipid peroxidation, Huh-7 cells (20000 cells per well) were plated in clear flat bottom 96 well plates in 200 µL DMEM and incubated for 24 h. After incubation, cells were treated for 90 minutes 0.02% DMSO, 20 µM of IITK4101 and IITK4102. After treatment, cells were stained with 5 µM of C11-BODIPY (Thermo) to cells in 200 µL of DMEM media for 60 minutes.in the incubator. After staining, cells were washed with 100 µL of 1X PBS for twice, cells were observed under microscope and images were captured under fluorescent microscope (Bio-Rad ZOE fluorescent cell imager) red channel (λ_ex_ −556 nm; λ_em_ −615 nm) for normal C11-BODIPY images and in the green channel (λ_ex_ 480 nm; λ_em_ 517 nm). For oxidised C11-BODIPY images. Image colocalizations and the fluorescence intensity quantifications were performed using ImageJ software. Bar graphs were plotted by using the Graphpad Prism 9.0 software. The data provided were representative of three or four independent experiments. The data provided were representative of three or four independent experiments.

### Organelle trackers and lipi green fluorescence imaging

For co-staining experiments, 20000 Huh-7 cells per well were plated in clear flat bottom 96 well plates in 200 µL DMEM and incubated for 24 h. After incubation, cells were treated for 90 minutes 0.02% DMSO, 15 µM of IITK4101 and IITK4102. After treatment, cells were fixed with 4% formaldehyde for 15 min and permeabilized with 0.2% tween 20 for 10 min. Then they were stained with 1 µM of lipi green, 1 µM of lysotracker (Thermo) and endoplasmic reticulum tracker (Thermo), 4 µM of mitotracker (Thermo) to cells separately in 100 µL of 1x PBS. Cells were stained for 60 min with lyso / ER trackers and overnight with mito tracker. Post staining, cells were washed for thrice with ice cold 1X PBS and images were captured under fluorescent microscope (Bio-Rad ZOE fluorescent cell imager) red channel (λ_ex_ - 556 nm; λ_em_ −615 nm) for lyso tracker, blue channel (λ_ex_ −355 nm; λ_em_ −433 nm) for mito and ER trackers and green channel (λ_ex_ 480 nm; λ_em_ 517 nm) for lipi green. Images colocalizations and fluorescence intensity quantifications were performed using ImageJ software. Bar graphs were plotted by using the Graphpad Prism 9.0 software. The data provided were representative of three or four independent experiments. The data provided were representative of three or four independent experiments.

### General protocol of Resazurin-cell viability assay

Cells were cultured in T75 flasks (Thermo) until they attain 70%-80% confluency. Cells were trypsinized with 4 mL of trypsin (0.25% Trypsin-EDTA - Gibco) and counted by using hemocytometer. In 384 well plate, 500 cells/well with 75 μL media were plated into 360 wells (Thermo). The remaining 24 wells were plated with only media. Plates were incubated for 24 hours. Then they were treated with test compounds in eight-points-three-fold dilutions, individually or in combination with other molecules for 72 hours. After 72 hours of incubation, the media was discarded from the wells and washed with 1X PBS. 0.02 mg/mL of resazurin (SRL) was made in DMEM media and 150 uL of resazurin was added to every well of 96 well plate. After addition of resazurin, cells were incubated for 6 h and fluorescence readings were recorded at λ_ex_ - 520 nm and λ_em_– 580 nm in multimode plate reader (BioTek-CYTATION 5-imaging reader). Percentage cell viability was calculated as ((fluorescence of treated cells - fluorescence of media control)/ (fluorescence of DMSO controls – fluorescence of media control)) x 100. Graphs were plotted by using non-linear regression curve fit programme of Graphpad Prism 9.0. EC_50_ was calculated using Graphpad Prism software. The data provided were representative of three or four independent experiments.

### Concentrations of compounds used for cell viability assay

For the dose dependent evaluation, 10 µM to 0.0045 µM of IITK4101, IITK4102 in triplicate were used in Huh-7 cells. 100 µM to 0.045 µM of IITK4101, IITK4102 were used in U2OS and HEK293 cells. For the combination cell viability assessment, 0.1 µM and 1 µM of IITK4101/IIT4102 along with 100 µM of H_2_O_2_, 30 µM of *^t^*BuOOH, 500 µM of NAC, 50 µM of *α*-tocopherol, 500 µM of GSH, 10 µM of Qvd-OPh (Adooq Biosciences), 10 µM of Necrostatin-1 (Adooq Biosciences), 100 nM of Erastin (Adooq Biosciences), 10 nM of RSL-3 (Adooq Biosciences), 100 nM of Liproxstatin-1 (AK Scientific), 10 µM of Ferrostatin (AK Scientific) and 30 nM of Fin56 (AK Scientific) were used in triplicate. Further combination cell viability assay was performed by using 10 µM to 0.0045 µM of IITK4101, IITK4102 along with 50 nM of Rapamyicin (sigma) and 10 nM of Bafilomycin A (sigma) in triplicate. For every experiment, 0.2% DMSO was used as negative control and 500 nM of Doxorubicin (sigma) and 5 µM of cisplatin (Alfa Aesar) were used as positive controls.

### Wound healing/scratch assay

For scratch assay, 0.1 million Huh-7 cells/well were plated in clear bottom 24 well plates (NEST) and incubated for 24 hours. After incubation, scratches were made on the bottom surface of plates, washed cells with 300 µL of DMEM media for thrice. Cells were treated 2 µM of IITK4101and IITK4102, for 72 hours. The scratches were monitored, and the images were captured in multimode plate reader (BioTek-CYTATION 5-imaging reader) at 0 h, 24 h, 48 h, 72 h. The percentage scratch area migrated by cells was calculated by using ImageJ software and the bar graphs were plotted by using Graphpad Prism 9.0 software. The data provided were representative of three or four independent experiments.

### Colony forming assay

For colony forming assay, 1500 Huh-7 cells/well were plated in clear bottom 24 well plates and incubated for 14 days by changing DMEM media for every 4 days. After incubation, colonies were treated with seven-point-three-fold dilution series of IITK4101 and IITK4102 with concentrations ranging from 20 µM to 0.027 µM in triplicate for 72 hours. 0.2% DMSO was served as negative control. Post treatment, colonies were washed with 300 µL of 1x PBS for twice and they were fixed with 4% formaldehyde for 30 min at room temperature. Then the colonies were stained with 500 µL of 0.5% crystal violet (SRL) in 1x PBS at room temperature in dark for 30 min. Then the dye was gently removed, and the colonies were washed five times with distilled water to remove the excess dye. The plates were dried at room temperature overnight and the images of whole plate were captured. Then the crystal violet in the colonies was dissolved by adding 500 µL of 20% methanol in water for 60 min on rocker and the absorbance was recorded at 570 nm.in multimode plate reader. Percentage viable colonies was calculated as ((absorbance of treated cells - absorbance of media control)/ (absorbance of DMSO controls – absorbance of media control)) x 100. Graphs were plotted by using non-linear regression curve fit programme of Graphpad Prism 9.0. The data provided were representative of three or four independent experiments.

### Immunoblotting

For immuno blotting, 1M cells/well (Huh-7) were plated in 6 well plate (NEST) and were incubated for 24 hours. Post 24 hours of incubation, cells were treated for 2 and 4 h with 0.02% DMSO, 1 mL of EBSS, 20 µM of IITK4101 and IITK4102, 100 nM of Bafilomycin A, Baf + IITK4101, Baf + IITK4102, Baf + EBSS. Post treatment, media was removed, cells were washed with 2 mL of ice cold 1X PBS. 60 µL of 1X protease phosphatase inhibitor (Thermo Fischer) was added per well and cells were scrapped. After scraping, cells were lysed by using probe sonicator and cell lysate was collected after centrifugation at 20,000g for 45 minutes at 4 °C. Protein was quantified by Bradford reagent (SRL) and BSA (SRL) standard. 20 µg protein was resolved on a 4-12% polyacrylamide gel for 90 minutes at 130 V. The gel was transferred to PVDF membrane (bio-rad) for 12 hours at 40 V, 4 °C by using transfer buffer (1.452g Tris-base, 7.2g glycine in 800 mL distilled H2O+200 mL of 100% methanol). The membrane was washed for 10 minutes with distilled water. The membrane was cut according to the molecular weight of the desired protein. The membranes were blocked for 1 hour with 5% skimmed milk (SRL) in 1X TBST (50 mL of 10x TBST (12.1g tris-base, 40g glycine in 500 mL dH2O at pH-7.6) + 450 mL dH2O + 250 µL Tween 20 (SRL). After blocking, the membranes were washed thrice every 10 minutes with 1X TBST. Then the membranes were incubated with primary antibodies in 1% skimmed milk with 1X TBST, i.e., GAPDH (Anti-Rabbit-1:2500 dilution-Abgenex), LC-III (Anti-Rabbit-1 to 1000 dilution), ATGL (Anti-Mouse antibody at 1 to 1000 dilution - Cat No. SC365278, gifted by Dr. Suresh Kumar of IIT Kanpur) for overnight at 4 °C. After incubation with primary antibody, the membranes were washed for thrice every 10 minutes with 1X TBST. Then the membranes were incubated with anti-rabbit secondary antibody (1:5000 dilution) or Anti-mouse IgG HRP linked antibody (at 1 to 2000 dilutuon, Cat No. 7076, Cell signalling and technology (CST)) in 1% skimmed milk for 1 hour at room temperature. After incubation, the membranes were washed thrice every 10 minutes with 1X TBST. Then blots were developed by using 250 mM luminol, 90 mM of *p*-Coumaric acid with 50% H_2_O_2_. The blots were visualized by using chemidoc instrument’s chemiluminescence filter.

### Immunofluorescence imaging studies

For immunofluorescence experiments in Huh-7 cells, 20000 cells per well were plated in clear bottom 96 well plate and incubated for 24 h. They were treated in triplicate with DMSO, 15 μM of IITK4101/IITK4102, 500 μM of LLOMe (Sigma) for 90 minutes in the incubator. Post treatment, cells were washed with 200 μL of ice cold 1x PBS and fixed with 100 μL of 4% formaldehyde for 15 minutes at room temperature. The cells were permeabilized with 0.2% tween-20 for 10 min and blocked with 3% BSA in 1x PBS for 15 min at RT. Then cells were added with 50 µL of primary antibodies in 3% BSA in the following dilutions i.e.TFEB (Anti-rabbit-1:200 dilution), Alix (Anti-rabbit-1:600 dilution), LAMP2 (Anti-mouse-1:200 dilution), Galectin3 (Anti-mouse-1:200 dilution), LC-III (Anti-rabbit-1:400 dilution). TFEB antibody was co-stained with Hoechst 342 (1 µg/mL), Alix was co-stained with LAMP2 and Hoechst 342, LC-III was co-stained with Gal3 and Hoechst 342. The cells were incubated with 1° antibodies for overnight at 4 °C. After incubation, they were washed for twice with 100 µL of 1x PBS and incubated with the following secondary antibodies for 1 hour at room temperature in 3% BSA. The secondary antibodies used were Alexa 568 (Anti-rabbit) for TFEB, Alix, LC-III and Alexa 488 (Anti-mouse) for LAMP2 and Gal3. The cells were washed twice with 100 µL of 1x PBS. The images were captured, and the fluorescence was recorded by using Thermo Scientific Multiskan plate reader. The co-localized images and readings were directly obtained from the plate reader. The bar graphs were plotted using GraphPad Prism 9.0 software.

## Acknowledgments

We thank Dr. N. Mohan, BSBE, IIT Kanpur for gifting cell lines (Huh-7 and U2OS). This work was supported by Department of Biotechnology, DBT-IYBA grant (BT/12/IYBA/2019/07), Extramural Research grant from Council of Scientific and Industrial Research (CSIR), India (02(366)/19/EMR-II), CRG from DST-Science and Engineering Research Board (SERB, CRG/2021/004787), Indian Council for Medical Research (ICMR, AMR/ADHOC/296/2022-ECD-II), SERB (CRG/2019/003058) and CSIR (01(3050)/21/EMR-II).

## Author contributions

D.A. and S.K.V. conceived the idea. D.A., S.K., and S.K.V., designed the experiments. S.K.V., and D.A., executed high-content imaging, quantification and analysis of LDS. S.K.V S.D., and D.A., conducted fluorescence imaging, quantification, cell viability experiments, western blotting and mass spectral lipidomics analysis. S.K.V., D.Ajnar., and S.K., performed transfection experiments, immunofluorescence imaging and quantification. S.S., R.K., A.D and D.A., are acknowledged for synthesis and characterization.

---

**Extended data figure 1:**
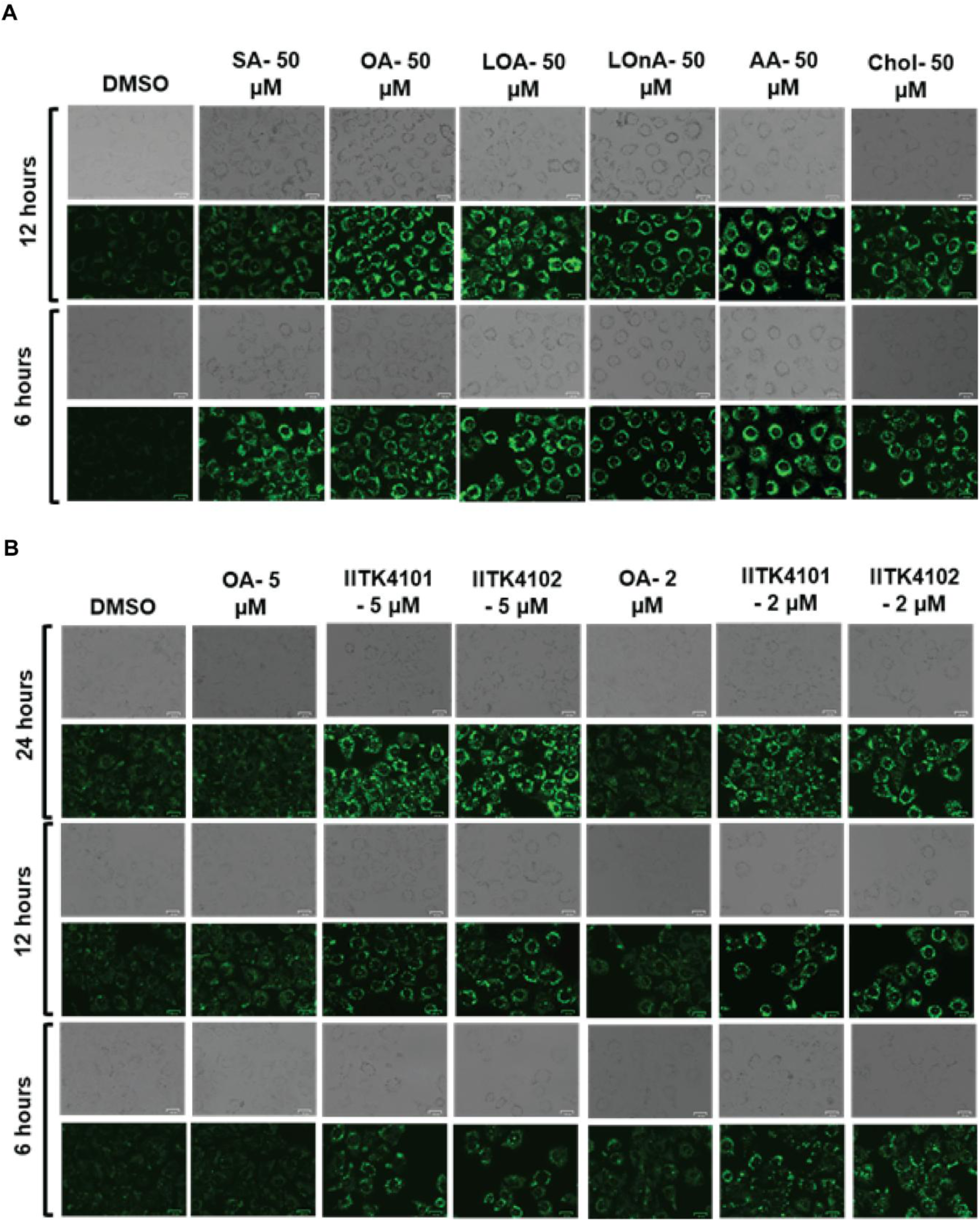
A, BRepresentative images of Huh-7 cells stained with lipi green. (A) Treatment conditions: DMSO (0.2%) and the mentioned fatty acids (50 µM) for 6 h and 12 h. (B). OA, IITK4101 and IITK4102 at 2 and 5 µm doses for 6, 12, and 24 h.

**Extended data figure 2:**
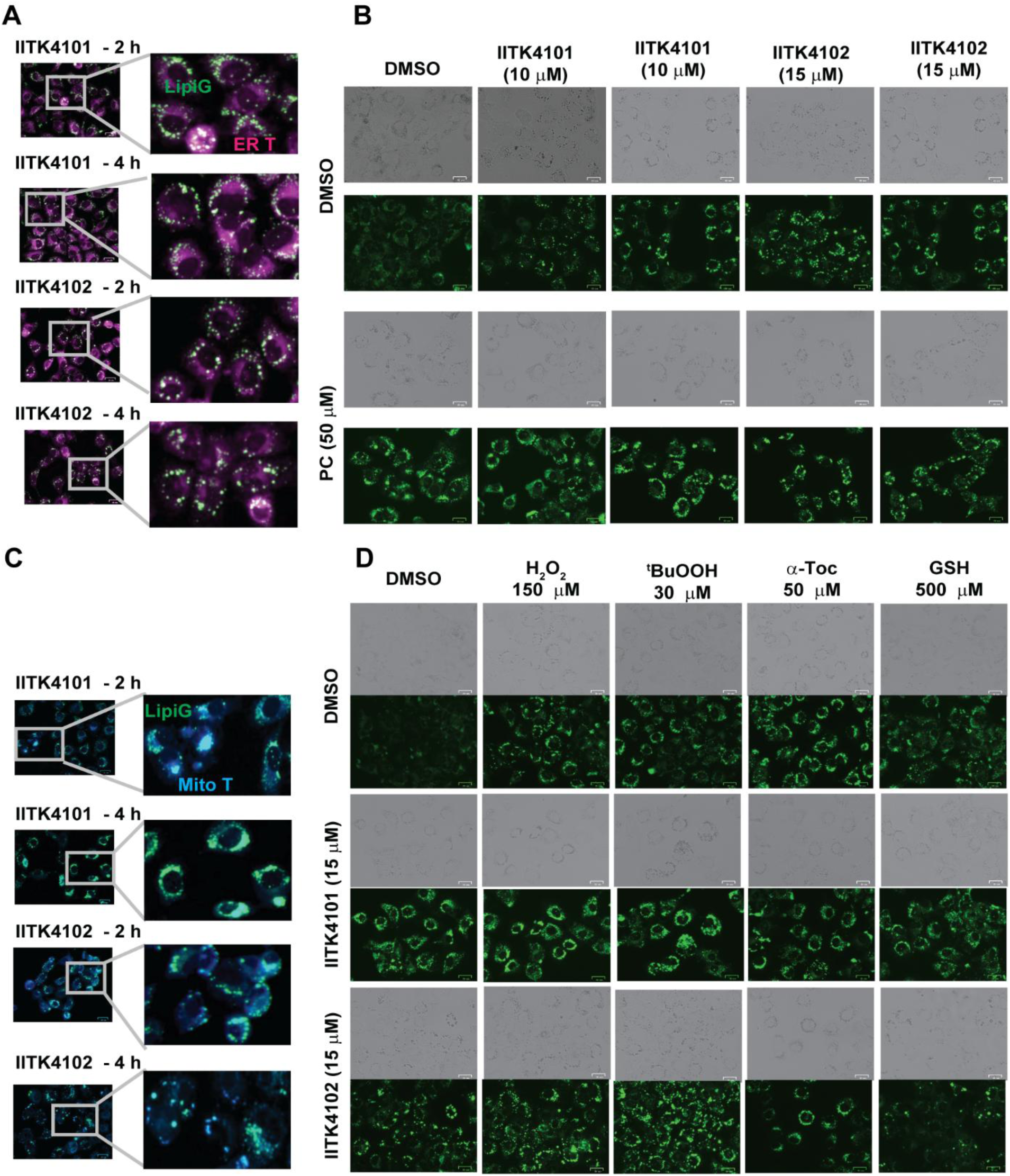
(A-D) Representative images of Huh-7 cells (A) co-stained with Endoplasmic reticulum tracker and lipi green after treatment with IITK4101 and IITK4102 (15 µM) for 2 h and 4 h, (B) %), phosphatidylcholine (50 µM), IITK4101 and IITK4102 (10 µM and 15 µM) and their combinations, (C) co-stained with Mito tracker and lipi green after treatment with IITK4101 and IITK4102 (15 µM) for 2 h and 4 h, and (D) with DMSO (0.2%), H_2_O_2_-150 µM, *^t^*BuOOH-30 µM, α-toc-50 µM, GSH-500 µM IITK4101/IITK4102 (15 µM) and their combinations for 3 h

**Extended data figure 3:**
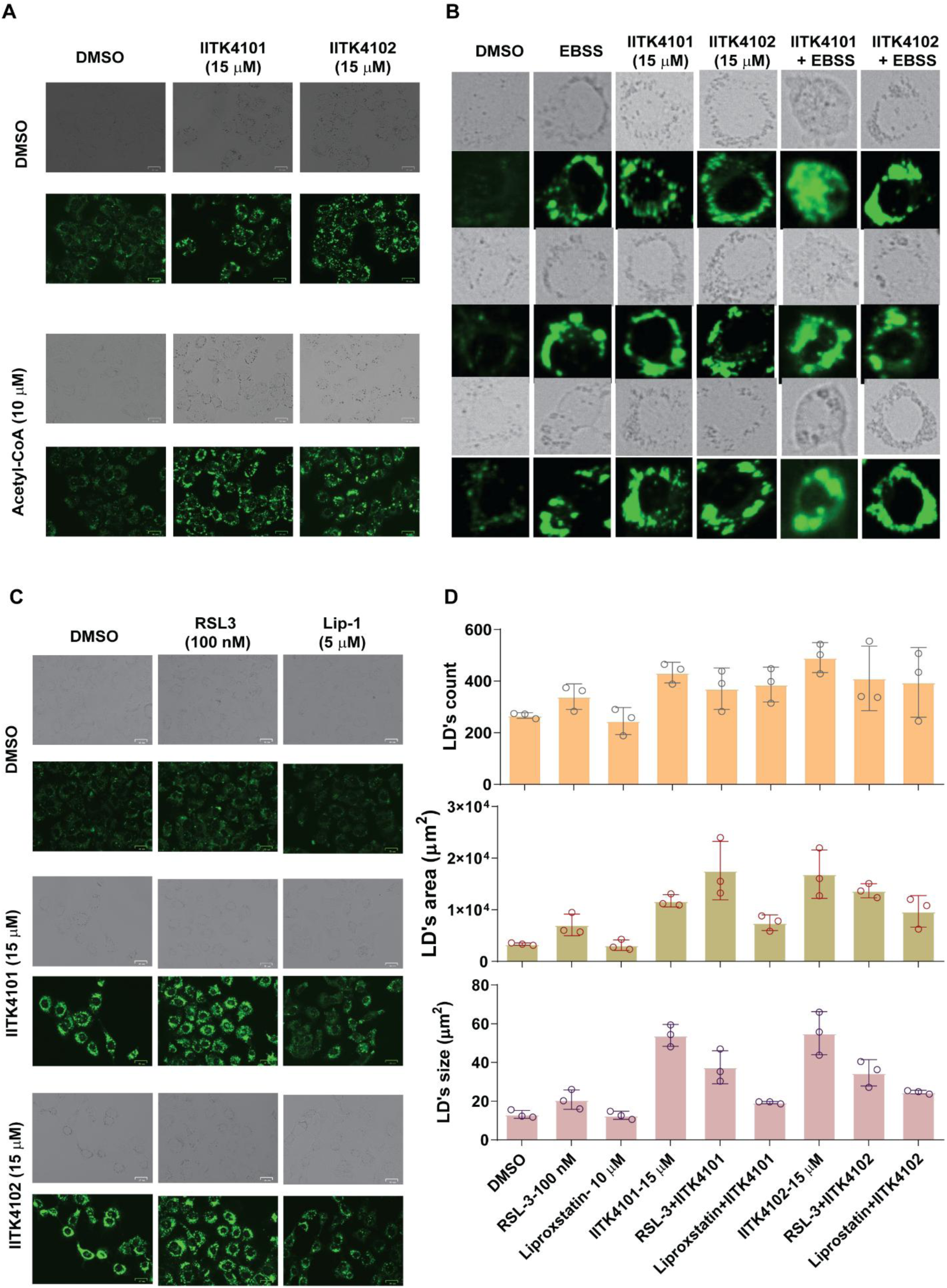
(A-D). Representative images of Huh-7 cells stained with lipi green after treatment with (A) DMSO (0.2%), Acetyl CoA (15 µM) IITK4101/IITK4102 (15 µM) and their combinations for 3 h, (B) DMSO (0.2%), EBSS (50 µL), IITK4101/IITK4102 (5 µM) and their combinations for 2 h, 4 h, 6 h. (C) DMSO (0.2%), RSL-3 (100 nM), Liproxstatin (5 µM), IITK4101/IITK4102 (15 µM) and their combinations for 3 h and (D) the quantification data of lipid droplets size, count and area.

**Extended data figure 4:**
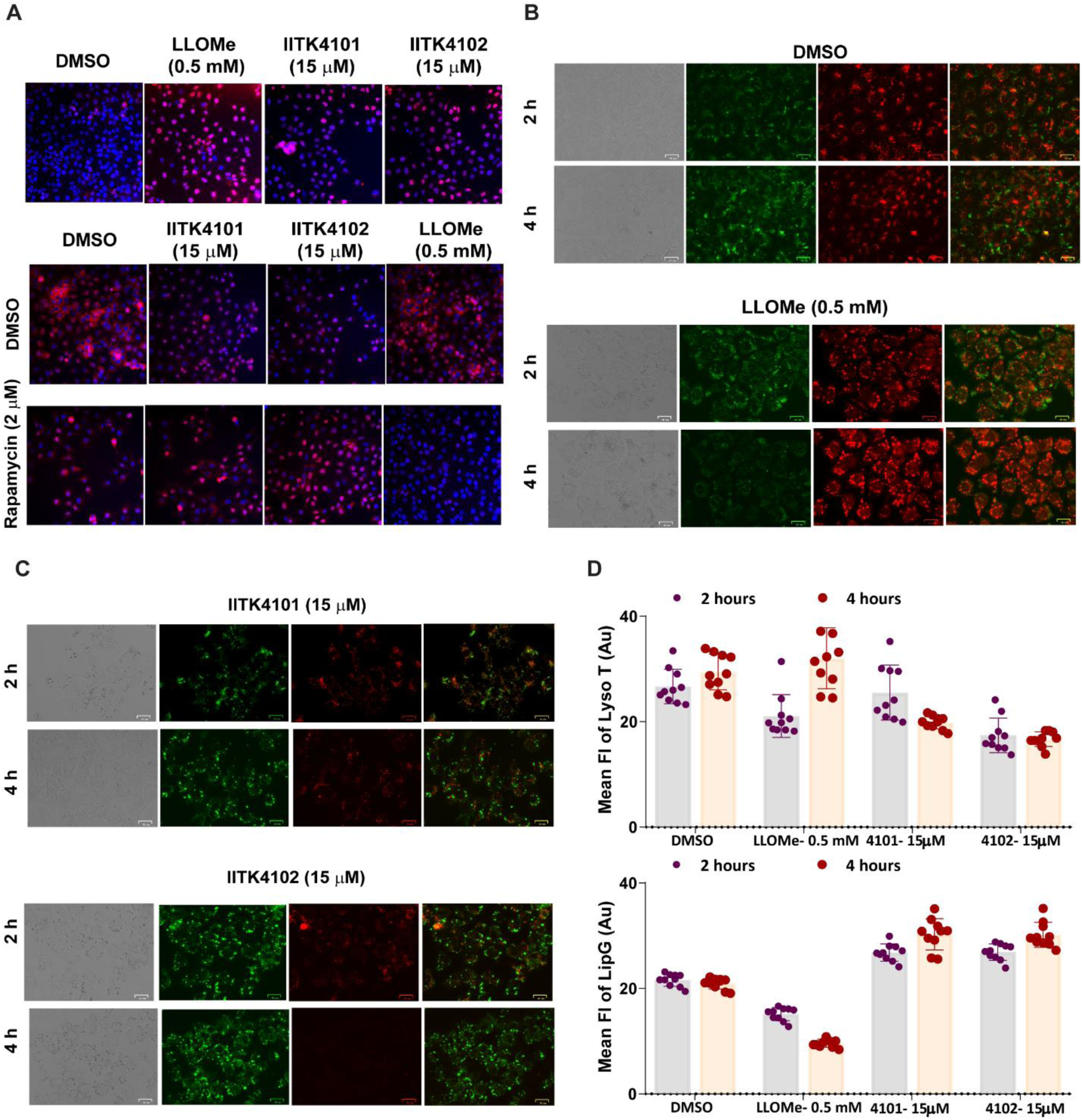
(A-D) Representative images of Huh-7 cells (A) co-stained with TFEB antibody and Hoechst after treatment with DMSO (0.2%), LLOMe (0.5 mM), IITK4101/IITK4102 (15 µM) for 3 h (A), in combination with Rapamycin (2 µM), (B) co-stained with Lyso tracker and lipi green after treatment with DMSO (0.2%) and LLOMe, (C) IITK4101 and IITK4102 (15 µM), for 2 h and 4 h, and (D) the quantified mean fluorescence intensities of lipi green and lyso tracker in the indicated treatment conditions.

## References

1 Olzmann, J. A. & Carvalho, P. Dynamics and functions of lipid droplets. Nature reviews. Molecular cell biology 20, 137–155, doi:10.1038/s41580-018-0085-z (2019).

2 Thiam, A. R., Farese Jr, R. V. & Walther, T. C. The biophysics and cell biology of lipid droplets. Nature reviews. Molecular cell biology 14, 775–786, doi:10.1038/nrm3699 (2013).

3 Londos, C., Brasaemle, D. L., Schultz, C. J., Segrest, J. P. & Kimmel, A. R. Perilipins, ADRP, and other proteins that associate with intracellular neutral lipid droplets in animal cells. Seminars in cell & developmental biology 10, 51–58, doi:10.1006/scdb.1998.0275 (1999).

4 Andersson, L. et al. PLD1 and ERK2 regulate cytosolic lipid droplet formation. J. Cell Sci. 119, 2246–2257, doi:10.1242/jcs.02941 %J Journal of Cell Science (2006).

5 Harayama, T. & Riezman, H. Understanding the diversity of membrane lipid composition. Nature reviews. Molecular cell biology 19, 281–296, doi:10.1038/nrm.2017.138 (2018).

6 Brown, D. A. Lipid droplets: proteins floating on a pool of fat. Current biology: CB 11, R446–449, doi:10.1016/s0960-9822(01)00257-3 (2001).

7 Mejhert, N. et al. The Lipid Droplet Knowledge Portal: A resource for systematic analyses of lipid droplet biology. Dev. Cell 57, 387–397.e384, 10.1016/j.devcel.2022.01.003 (2022).

8 Tauchi-Sato, K., Ozeki, S., Houjou, T., Taguchi, R. & Fujimoto, T. The Surface of Lipid Droplets Is a Phospholipid Monolayer with a Unique Fatty Acid Composition*. The Journal of biological chemistry 277, 44507–44512, 10.1074/jbc.M207712200 (2002).

9 Bersuker, K. et al. A Proximity Labeling Strategy Provides Insights into the Composition and Dynamics of Lipid Droplet Proteomes. Dev. Cell 44, 97–112.e117, 10.1016/j.devcel.2017.11.020 (2018).

10 Zadoorian, A., Du, X. & Yang, H. Lipid droplet biogenesis and functions in health and disease. Nat. Rev. Endocrinol. 19, 443–459, doi:10.1038/s41574-023-00845-0 (2023).

11 Herms, A. et al. Cell-to-cell heterogeneity in lipid droplets suggests a mechanism to reduce lipotoxicity. Current biology: CB 23, 1489–1496, doi:10.1016/j.cub.2013.06.032 (2013).

12 Nguyen, T. B. et al. DGAT1-Dependent Lipid Droplet Biogenesis Protects Mitochondrial Function during Starvation-Induced Autophagy. Dev. Cell 42, 9–21.e25, doi:10.1016/j.devcel.2017.06.003 (2017).

13 Herms, A. et al. AMPK activation promotes lipid droplet dispersion on detyrosinated microtubules to increase mitochondrial fatty acid oxidation. Nature communications 6, 7176, doi:10.1038/ncomms8176 (2015).

14 Valm, A. M. et al. Applying systems-level spectral imaging and analysis to reveal the organelle interactome. Nature 546, 162–167, doi:10.1038/nature22369 (2017).

15 Talari, N. K. et al. Lipid-droplet associated mitochondria promote fatty-acid oxidation through a distinct bioenergetic pattern in male Wistar rats. Nature communications 14, 766, doi:10.1038/s41467-023-36432-0 (2023).

16 Benador, I. Y. et al. Mitochondria Bound to Lipid Droplets Have Unique Bioenergetics, Composition, and Dynamics that Support Lipid Droplet Expansion. Cell metabolism 27, 869–885.e866, doi:10.1016/j.cmet.2018.03.003 (2018).

17 Nakajima, S., Gotoh, M., Fukasawa, K., Murakami-Murofushi, K. & Kunugi, H. Oleic acid is a potent inducer for lipid droplet accumulation through its esterification to glycerol by diacylglycerol acyltransferase in primary cortical astrocytes. Brain Res. 1725, 146484, 10.1016/j.brainres.2019.146484 (2019).

18 Castillo, H. B., Shuster, S. O., Tarekegn, L. H. & Davis, C. M. Oleic acid differentially affects lipid droplet storage of de novo synthesized lipids in hepatocytes and adipocytes. Chem. Commun. 60, 3138–3141, doi:10.1039/D3CC04829B (2024).

19 Rohwedder, A., Zhang, Q., Rudge, S. A. & Wakelam, M. J. O. Lipid droplet formation in response to oleic acid in Huh-7 cells is mediated by the fatty acid receptor FFAR4. J. Cell Sci. 127, 3104–3115, doi:10.1242/jcs.145854 (2014).

20 Carro, M., Buschiazzo, J., Ríos, G. L., Oresti, G. M. & Alberio, R. H. Linoleic acid stimulates neutral lipid accumulation in lipid droplets of maturing bovine oocytes. Theriogenology 79, 687–694, 10.1016/j.theriogenology.2012.11.025 (2013).

21 Zimmermann, R. et al. Fat Mobilization in Adipose Tissue Is Promoted by Adipose Triglyceride Lipase. Science (New York, N.Y.) 306, 1383–1386, doi:10.1126/science.1100747 (2004).

22 Haemmerle, G. et al. Defective Lipolysis and Altered Energy Metabolism in Mice Lacking Adipose Triglyceride Lipase. Science (New York, N.Y.) 312, 734–737, doi:10.1126/science.1123965 (2006).

23 Lange, M. & Olzmann, J. A. Ending on a sour note: Lipids orchestrate ferroptosis in cancer. Cell metabolism 33, 1507–1509, 10.1016/j.cmet.2021.07.011 (2021).

24 Lange, M., Wagner, P. V. & Fedorova, M. Lipid composition dictates the rate of lipid peroxidation in artificial lipid droplets. Free Radical Res. 55, 469–480, doi:10.1080/10715762.2021.1898603 (2021).

25 Ferrada, L., Barahona, M. J., Vera, M., Stockwell, B. R. & Nualart, F. Dehydroascorbic acid sensitizes cancer cells to system xc-inhibition-induced ferroptosis by promoting lipid droplet peroxidation. Cell Death Dis. 14, 637, doi:10.1038/s41419-023-06153-9 (2023).

26 Lange, M. et al. FSP1-mediated lipid droplet quality control prevents neutral lipid peroxidation and ferroptosis. bioRxiv, 2025.2001.2006.631537, doi:10.1101/2025.01.06.631537 %J bioRxiv (2025).

27 Tatenaka, Y. et al. Monitoring Lipid Droplet Dynamics in Living Cells by Using Fluorescent Probes. Biochemistry 58, 499–503, doi:10.1021/acs.biochem.8b01071 (2019).

28 Smirnova, E. et al. ATGL has a key role in lipid droplet/adiposome degradation in mammalian cells. EMBO reports 7, 106–113, doi:10.1038/sj.embor.7400559 (2006).

29 Haemmerle, G., Zimmermann, R. & Zechner, R. Letting lipids go: hormone-sensitive lipase. Current opinion in lipidology 14, 289–297, doi:10.1097/00041433-200306000-00009 (2003).

30 Cabruja, M. et al. In-depth triacylglycerol profiling using MS3 Q-Trap mass spectrometry. Anal. Chim. Acta 1184, 339023, 10.1016/j.aca.2021.339023 (2021).

31 Hartler, J., Köfeler, H. C., Trötzmüller, M., Thallinger, G. G. & Spener, F. Assessment of lipidomic species in hepatocyte lipid droplets from stressed mouse models. Sci. Data 1, 140051, doi:10.1038/sdata.2014.51 (2014).

32 Chitraju, C. et al. Lipidomic analysis of lipid droplets from murine hepatocytes reveals distinct signatures for nutritional stress. J. Lipid Res. 53, 2141–2152, 10.1194/jlr.M028902 (2012).

33 Chitraju, C. et al. The impact of genetic stress by ATGL deficiency on the lipidome of lipid droplets from murine hepatocytes. J. Lipid Res. 54, 2185–2194, 10.1194/jlr.M037952 (2013).

34 Peterson, C. W. H., Deol, K. K., To, M. & Olzmann, J. A. Optimized protocol for the identification of lipid droplet proteomes using proximity labeling proteomics in cultured human cells. STAR Protocols 2, 100579, 10.1016/j.xpro.2021.100579 (2021).

35 Kennedy, E. P. & Weiss, S. B. The function of cytidine coenzymes in the biosynthesis of phospholipides. The Journal of biological chemistry 222, 193–214 (1956).

36 Krahmer, N. et al. Phosphatidylcholine synthesis for lipid droplet expansion is mediated by localized activation of CTP:phosphocholine cytidylyltransferase. Cell metabolism 14, 504–515, doi:10.1016/j.cmet.2011.07.013 (2011).

37 Vechalapu, S. K. et al. Redox modulator iron complexes trigger intrinsic apoptosis pathway in cancer cells. iScience 27, doi:10.1016/j.isci.2024.109899 (2024).

38 Dharmaraja, A. T. & Chakrapani, H. A Small Molecule for Controlled Generation of Reactive Oxygen Species (ROS). Org. Lett. 16, 398–401, doi:10.1021/ol403300a (2014).

39 Garcia-Irigoyen, O. et al. Enterocyte superoxide dismutase 2 deletion drives obesity. iScience 25, 103707, doi:10.1016/j.isci.2021.103707 (2022).

40 Dixon, Scott J. et al. Ferroptosis: An Iron-Dependent Form of Nonapoptotic Cell Death. Cell 149, 1060–1072, 10.1016/j.cell.2012.03.042 (2012).

41 Kassan, A. et al. Acyl-CoA synthetase 3 promotes lipid droplet biogenesis in ER microdomains. The Journal of cell biology 203, 985–1001, doi:10.1083/jcb.201305142 (2013).

42 Dharmaraja, A. T. Role of Reactive Oxygen Species (ROS) in Therapeutics and Drug Resistance in Cancer and Bacteria. J. Med. Chem. 60, 3221–3240, doi:10.1021/acs.jmedchem.6b01243 (2017).

43 Jin, Y., Tan, Y., Wu, J. & Ren, Z. Lipid droplets: a cellular organelle vital in cancer cells. Cell Death Discov. 9, 254, doi:10.1038/s41420-023-01493-z (2023).

44 Cruz, A. L. S., Barreto, E. d. A., Fazolini, N. P. B., Viola, J. P. B. & Bozza, P. T. Lipid droplets: platforms with multiple functions in cancer hallmarks. Cell Death Dis. 11, 105, doi:10.1038/s41419-020-2297-3 (2020).

45 Yang, Z. & Klionsky, D. J. Eaten alive: a history of macroautophagy. Nature cell biology 12, 814–822, doi:10.1038/ncb0910-814 (2010).

46 Mizushima, N. & Komatsu, M. Autophagy: renovation of cells and tissues. Cell 147, 728–741, doi:10.1016/j.cell.2011.10.026 (2011).

47 Ichimura, Y. et al. A ubiquitin-like system mediates protein lipidation. Nature 408, 488–492, doi:10.1038/35044114 (2000).

48 Kabeya, Y. et al. LC3, a mammalian homologue of yeast Apg8p, is localized in autophagosome membranes after processing. Embo J. 19, 5720–5728, doi:10.1093/emboj/19.21.5720 (2000).

49 Yamamoto, A. et al. Bafilomycin A1 Prevents Maturation of Autophagic Vacuoles by Inhibiting Fusion between Autophagosomes and Lysosomes in Rat Hepatoma Cell Line, H-4-II-E Cells. Cell Struct. Funct. 23, 33-42, doi:10.1247/csf.23.33 (1998).

50 Klionsky, D. J., Elazar, Z., Seglen, P. O. & Rubinsztein, D. C. Does bafilomycin A1 block the fusion of autophagosomes with lysosomes? Autophagy 4, 849–850, doi:10.4161/auto.6845 (2008).

51 Lapierre, L. R. et al. The TFEB orthologue HLH-30 regulates autophagy and modulates longevity in Caenorhabditis elegans. Nature communications 4, 2267, doi:10.1038/ncomms3267 (2013).

52 Sardiello, M. et al. A gene network regulating lysosomal biogenesis and function. Science (New York, N.Y.) 325, 473–477, doi:10.1126/science.1174447 (2009).

53 Settembre, C. & Ballabio, A. TFEB regulates autophagy: an integrated coordination of cellular degradation and recycling processes. Autophagy 7, 1379–1381, doi:10.4161/auto.7.11.17166 (2011).

54 Thiele, D. L. & Lipsky, P. E. Mechanism of L-leucyl-L-leucine methyl ester-mediated killing of cytotoxic lymphocytes: dependence on a lysosomal thiol protease, dipeptidyl peptidase I, that is enriched in these cells. Proceedings of the National Academy of Sciences of the United States of America 87, 83–87, doi:10.1073/pnas.87.1.83 (1990).

55 Radulovic, M. & Stenmark, H. Lysophagy prevents neurotoxic aggregate transmission. Proceedings of the National Academy of Sciences of the United States of America 121, e2321181121, doi:10.1073/pnas.2321181121 (2024).

56 Kumar, S. et al. Galectins and TRIMs directly interact and orchestrate autophagic response to endomembrane damage. Autophagy 13, 1086–1087, doi:10.1080/15548627.2017.1307487 (2017).

57 Nguyen, T. N. et al. Atg8 family LC3/GABARAP proteins are crucial for autophagosome-lysosome fusion but not autophagosome formation during PINK1/Parkin mitophagy and starvation. J Cell Biol. 215, 857–874, doi:10.1083/jcb.201607039 (2016).

58 Eriksson, I., Wäster, P. & Öllinger, K. Restoration of lysosomal function after damage is accompanied by recycling of lysosomal membrane proteins. Cell Death Dis. 11, 370, doi:10.1038/s41419-020-2527-8 (2020).

59 Skowyra, M. L., Schlesinger, P. H., Naismith, T. V. & Hanson, P. I. Triggered recruitment of ESCRT machinery promotes endolysosomal repair. Science (New York, N.Y.) 360, doi:10.1126/science.aar5078 (2018).

60 Radulovic, M. et al. ESCRT-mediated lysosome repair precedes lysophagy and promotes cell survival. Embo j. 37, doi:10.15252/embj.201899753 (2018).

61 Scheffer, L. L. et al. Mechanism of Ca²⁺-triggered ESCRT assembly and regulation of cell membrane repair. Nat Commun. 5, 5646, doi:10.1038/ncomms6646 (2014).

62 Jia, J. et al. Galectin-3 Coordinates a Cellular System for Lysosomal Repair and Removal. Developmental cell 52, 69–87.e68, doi:10.1016/j.devcel.2019.10.025 (2020).

63 Xu, S., Zhang, X. & Liu, P. Lipid droplet proteins and metabolic diseases. Biochim. Biophys. Acta Mol. Basis Dis. 1864, 1968–1983, doi:10.1016/j.bbadis.2017.07.019 (2018).

64 Greenberg, A. S. et al. The role of lipid droplets in metabolic disease in rodents and humans. J. Clin. Investig. 121, 2102–2110, doi:10.1172/JCI46069 (2011).

65 Martinez-Botas, J. et al. Absence of perilipin results in leanness and reverses obesity in Lepr(db/db) mice. Nat. Genet. 26, 474–479, doi:10.1038/82630 (2000).

